# Exploring metalloproteome remodeling in calprotectin-stressed *Acinetobacter baumannii* using chemoproteomics

**DOI:** 10.1101/2025.09.16.676631

**Authors:** Maximillian K. Osterberg, Daniel W. Bak, Claudia Andreini, Jeanette M. Critchlow, Jonathan C. Trinidad, Peter V. Cornish, Tae Akizuki, Walter J. Chazin, Eric P. Skaar, Eranthie Weerapana, David P. Giedroc

## Abstract

The growth of bacterial pathogens is limited by nutritional immunity, where the infected host deploys metal scavenging proteins to starve the pathogen of essential transition metals. An important transition metal-sequestering protein is the S100A8-S100A9 heterotetramer, calprotectin (CP). Prior work reveals that CP induces a significant Zn- and Fe-starvation response in the Gram-negative opportunistic pathogen, *Acinetobacter baumannii*, in liquid culture. Here, we employ a quantitative chemoproteomics platform to pinpoint changes in abundance-corrected cysteine reactivity, and by extension cellular metal occupancy in metalloenzymes, that occur when *A. baumannii* is challenged with physiological CP in liquid culture relative to an untreated WT control. Changes in protein abundance with CP stress reveal a pronounced Zn-limitation and Fe-starvation response and reciprocal regulation of three enzymes of central carbon metabolism, including aconitase. A majority of the 2645 quantifiable Cys-containing peptides that show an increase in abundance-corrected Cys reactivity (150) are derived from known Zn-, Fe- and Fe-S-cluster proteins, revealing a significant decrease in metal occupancy (undermetalation) across the proteome. Myriad cell processes are compromised by undermetalation of the metalloproteome, including enzymes that function in the TCA cycle and respiration, GTP metabolism, ribosome remodeling, tRNA charging, and proteostasis. A direct comparison of a strain lacking the candidate metallochaperone ZigA (Δ*zigA*) with the wild-type strain reveals that the loss of ZigA is effectively silent in this assay. We conclude that CP induces a widespread, negative impact on the metalation status of the metalloproteome that results in a significant nutrient limitation response.

## INTRODUCTION

The emergence of widespread antimicrobial resistance (AMR) to conventional and last-resort antibiotic therapies continues to complicate efforts to combat bacterial infectious disease in humans.^1–2^ One attractive approach to meet this challenge is to identify metabolic pathways in bacterial pathogens that become critical only during the course of an infection, in an adaptive often bacteriostatic response to host efforts to minimize dissemination from the initial site of colonization. This might ultimately lead to optimal deployment of myriad antimicrobial weapons by the host that are used to clear the invading pathogen. Targeting specific aspects of bacterial metallostasis,^3^ a term encompassing all processes that maintain the integrity of the metalloproteome upon challenge by metal limitation or metal toxicity,^4–6^ might be one such strategy to aid development of new antibacterial therapeutic strategies.^3,7^

An important antimicrobial response of the innate immune system toward extracellular pathogens is the deployment of transition metal-sequestering proteins that limit the availability of transition metals to the bacterium at sites of infection. This induces a metal starvation response in a process known as nutritional immunity.^4,8–9^ Originally discussed in the context of host nutrient Fe(III)-limitation mediated by lactoferrin, which binds Fe(III), and lipocalin-2, which binds Fe(III)-siderophore complexes, nutritional immunity against invading pathogens is now known to extend to other first-row *d*-block metals including Mn(II) and Zn(II) as well as Fe(II).^10–14^ The S100 sub-family of EF-hand Ca(II)-binding proteins have distinct transition-metal binding sites that provide them with defined roles in host nutritional immunity. The S100A8/S100A9 heterotetramer, calprotectin (CP),^15–16^ is a unique member of this sub-family, well characterized for its participation in the “tug-of-war” between pathogen and host for nutrient metals.^17^ CP harbors two pairs of high-affinity transition-metal coordination sites: One is a His_3_-Asp site that is common to other S100 proteins,^15,18–19^ while the other is an octahedral hexahistidine (His_6_) site found only in CP, which chelates Zn(II),^20^ Cu(II),^21^ Mn(II),^22^ Ni(II)^23^ and Fe(II).^24^ While the transition metal-sequestering role of CP is paramount to its function, evidence has emerged to suggest that CP induces a cell wall stress response that is independent of its role in metal sequestration.^25–26^ These findings largely derive from bacterial growth experiments in liquid culture in the presence of wild-type and mutant CP derivatives.

It is well-established that bacteria cultured in the presence of CP are starved for transition metals in a way that appears strongly dependent on the culture conditions and the organism itself.^15^ We have been interested in how the Gram-negative opportunistic and nosocomial pathogen, *Acinetobacter baumannii*, responds to CP-mediated stress.^27^ Early bacterial cell culture and metal add-back experiments revealed that CP induces Zn(II)-starvation in *A. baumannii* given the strong transcriptional up-regulation of the Zur (zinc-uptake repressor) regulon.^28–29^ Subsequent global transcriptomics and proteomics experiments revealed that CP drives a chronic Fe-starvation response superimposed on the Zn-limitation response.^27,30^ These studies also established that one of the metabolic responses to low Fe availability is the prioritization of flavin biosynthesis, and evidence was presented for a “work-around” strategy allowing cells to accumulate high levels of flavin required for electron transfer that might be compromised as a result of a decrease in Fe-containing ferredoxins.^30^ In addition, these studies hinted at a failure or prioritization of queuosine-tRNA biosynthesis, given the increased cellular abundance of the 6-carboxy-5,6,7,8-tetrahydropterin (6-CPH_4_) synthase (QueD2) which catalyzes the rate-determining step in this pathway; it was subsequently shown that CP treatment reduces flux through the queuosine-tRNA biosynthesis pathway.^31^ Queuosine is a hypermodified 7-deazaguanosine derivative^32–33^ found in the anticodon loop of four transfer RNAs (tRNA) and is upregulated under nutrient-limitation and metal-limitation stress responses in other bacteria and organisms.^34–35^ The Q substitution is known to impact tRNA structure and function in a way that can affect the rate and/or fidelity of translation and ultimately the composition of the proteome.^36^

The set-point model for transition metal homeostasis predicts that CP impacts bacterial metabolism by impairing the ability of the cellular milieu to supply metalloproteins with their cognate metal or metal cofactor.^37–38^ This model predicts widespread undermetalation of some or all the metalloproteome,^3,37^ as has been demonstrated for selected metalloenzymes in a small number of organisms.^23,39–40^ The extent to which the metalloproteome is globally impacted in this way is currently unknown. Here, we employ a quantitative chemoproteomics platform to elucidate changes in abundance-corrected cysteine (Cys) reactivity that results when *A. baumannii* is exposed to physiological CP in liquid culture relative to an untreated control. Metal-liganding cysteines in a metalloprotein become more reactive toward an electrophile upon metal dissociation, since metal-coordination attenuates the nucleophilicity of that cysteine.^41–44^ Thiolate ligands are ubiquitous components of Fe-S cluster-containing proteins and a subset of Zn-metalloprotein sites, and as a result changes in the reactivity of these cysteines can act as a proxy for cellular metallocofactor occupancy.^45^ Strong support for this hypothesis derives from pioneering studies of Fe-S protein biogenesis in *Escherichia coli,*^46–48^ and examination of the effects of the thiophilic metal-chelating prodrug holomycin in *E. coli.*^49^

We reasoned that this chemoproteomic workflow could be used to globally probe metallocofactor occupancy in CP-treated *A. baumannii*, leveraging the finding that CP induces a Zn- and Fe-limitation regulatory response in this organism.^30–31^ Using an isobaric mass spectrometry-based tagging strategy to quantify protein abundance, we show that pairs of enzymes involved in central carbon metabolism show reciprocal changes in protein abundance. These include two pairs of TCA cycle enzymes, aconitase^50^ and fumarase,^49^ a Zn- vs. Fe-type alcohol dehydrogenase, and the ribosomal proteins L31 and the low-zinc paralog L31B,^51–52^ these findings consistent with a classical metal-sparing response.^53–54^ In total, 29 Zn and Fe-S cluster proteins were found to exhibit a statistically significant ≥2-fold increase in the reactivity of metal-coordinating cysteines, including 22 Zn-metalloenzymes and 7 Fe-S cluster proteins.

Undermetalation of Fe-S proteins negatively impacts their cell abundance, potentially consistent with increased turnover and more globale Fe sparing, in contrast to the vast majority of undermetalated zinc enzymes, which display minimal changes in abundance. These undermetalated enzymes function in energy metabolism and respiration, GTP metabolism, ribosome remodeling and tRNA charging, RNA metabolism and proteostasis. In addition, ribosomes from CP-stressed cells exhibit classical hallmarks of ribosome hibernation and nutrient limitation.^55^ These findings are broadly recapitulated in a strain lacking the candidate Zur-regulated metallochaperone, ZigA,^27,56–57^ leaving unknown any potential metalloenzyme client for this GTPase.

## RESULTS

### Experimental Strategy

In this work, we employed a 6-plex amine-reactive tandem mass-tag (TMT)-labeling approach coupled with desthiobiotin iodoacetamide (DBIA) Cys-labeling and streptavidin-agarose biotin enrichment for sequential quantification of protein abundance and abundance-corrected cysteine-reactivity changes in CP-stressed vs. unstressed *A. baumannii* ATCC 17978-VU cells (Figure 1). Biological duplicate bacterial cell lysates containing both soluble and membrane-associated proteins were obtained for untreated wild-type (WT), CP (300 µg/mL)-stressed WT and CP-stressed Δ*zigA* cells (six samples total) and subjected to DBIA labeling and protein isolation, digested with trypsin and subjected to 6-plex TMT labeling using the scheme indicated (Figure 1). The six samples were combined together and one part of the combined sample (≈25% of the total; in technical duplicate) was subjected directly to isobaric mass-spectrometry analysis to quantify relative protein abundance across each TMT channel. This experimental workflow thus provides a quantitative read-out of the relative changes in protein abundance in three pair-wise comparisons (measured across four replicate channels per biological condition): 1) WT vs. WT+CP; 2) WT vs. Δ*zigA* +CP, and 3) WT + CP vs. Δ*zigA* + CP in an approach analogous to label-free proteomics experiments published earlier.^30^ This concentration of CP is sufficient to significantly sequester Zn, Fe(II), and Mn from the media (Figure S1A), yet induces only a significant ≈2-fold decrease in total cell-associated Zn as measured by inductively coupled plasma-mass spectrometry (ICP-MS), with little corresponding change in total cell-associated Fe despite these cells being significantly Fe-starved^30^ (Figure S1B).

**Figure 1.**
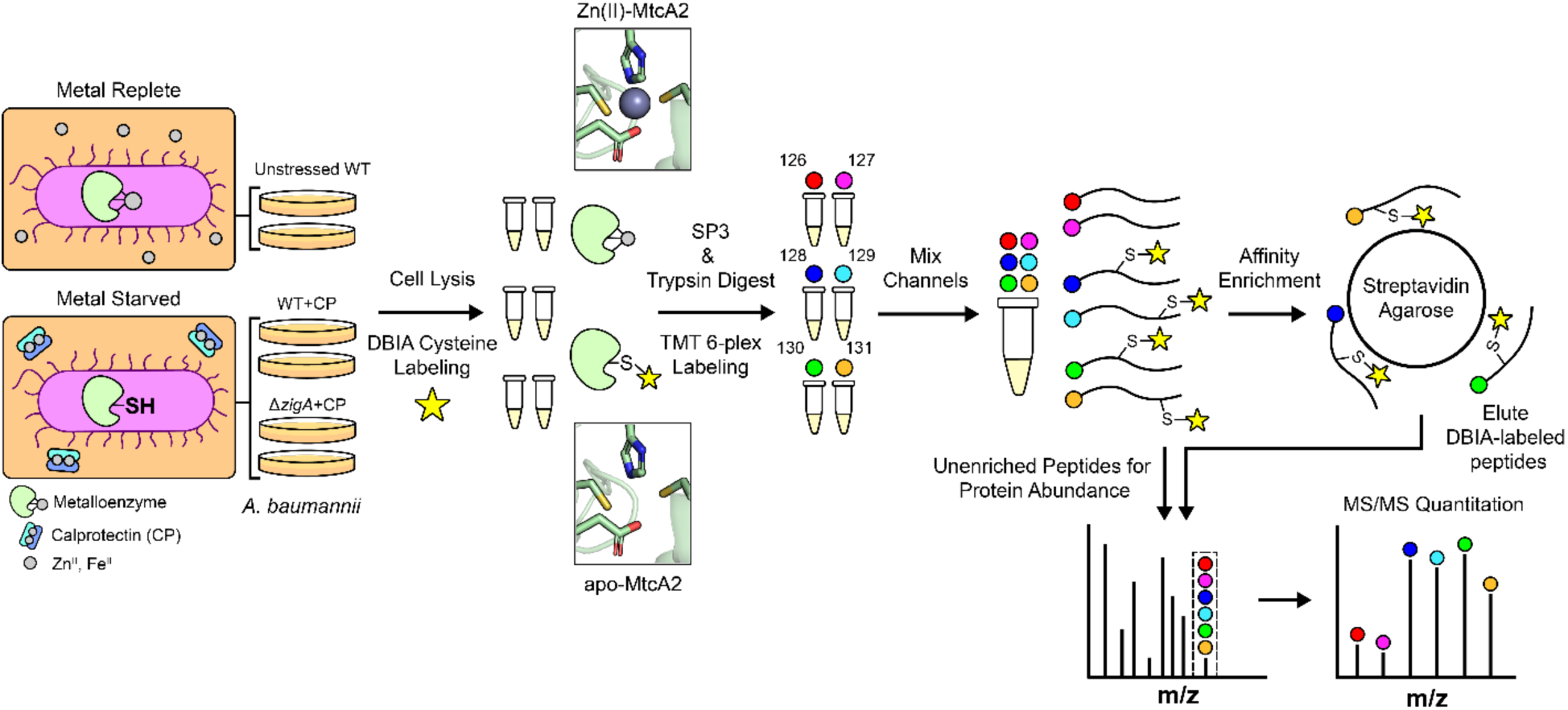
Schematic representation of the workflow employed in this study. Briefly, biological replicates of each of three bacterial cultures were subjected to cell lysis and thiol labeling, protein isolation and digestion, and derivatization with one of six TMT reagents, as indicated. Samples were mixed at 1:1:1:1:1:1 and this combined sample subjected to either protein abundance quantitation or streptavidin enrichment and cysteine reactivity quantitation by LC-MS/MS. The insets show ribbon representations of Zn(II) bound active site of β-carbonic anhydrase, MtcA2 (*upper*) and apo (metal-free) MtcA2 (*lower*) that might be present in an CP-treated sample (designated undermetallated). See text for additional details.

The remainder of the combined sample (75% of the total; in technical triplicate) was subjected to streptavidin enrichment and elution, with the relative abundance of individual eluted peptides again quantified by reporter-ion intensity. This provides relative quantitation of the abundance of Cys-containing peptides across the same three pair-wise comparisons (measured across six replicate channels per biological condition). Normalizing these cysteine-reactivity changes against the changes in protein abundance, a net change in Cys reactivity is obtained as proxy for the relative change in fractional metal occupancy for a given metalloprotein.^44,47,49^ Only those changes in net Cys reactivity greater than 2-fold (log_2_=±1) with a χ^2^ cut-off of 9.0 were generally considered of primary significance here, although in selected instances we discuss changes with slightly lower values (log_2_=±0.64; χ^2^≥9.0).

We used MetalPredator,^58^ the metalPDB database^59^ and related tools with manual curation to estimate the number of Zn metalloproteins and Fe-S cluster proteins in the *A. baumannii* ATCC 17978-VU proteome to provide context for the findings reported here. We predict the presence of 225 zinc metalloproteins (comparable to 215 predicted previously^30^), of which 40% (87) contain at least one cysteine in the first coordination shell (Document S1). Of these 87 Zn proteins, 6 contain a single Cys, 11 contain two Cys, 22 contain three Cys, and 50 are tetrathiolate coordination sites that are known or anticipated to bind Zn, but if not previously characterized, could also potentially coordinate a 4Fe-4S cluster.^60^ Likewise, we estimate the presence of 92 Fe-S-cluster proteins in *A. baumanni* ATCC 17978-VU (Document S1). We note that *A. baumannii*, like *Pseudomonas aeruginosa*, contains the minimal complement of ISC enzymes required to assemble both [2Fe-2S] and [4Fe-4S] proteins,^61^ including the cysteine desulfurase IscS, the scaffolding protein IscU, the Fe-S cluster carriers IscA, ErpA, and NfuA,^62^ as well as two copies of the regulatory protein IscR.^63^ There is no “stress-responsive” SUF Fe-S cluster biogenesis system in *A. baumannii* like that found in *E. coli.*^48^ In all, we were able to detect 147 (65%) known or predicted Zn enzymes and 59 (64%) known or predicted Fe-S cluster proteins in the proteome (Figure 2A). These percentages represent excellent coverage, since they are slightly higher than the fraction of the total *A. baumannii* proteome detected in these experiments (1966 proteins or 52%); we note however that there may be unidentified Zn and Fe-S cluster proteins that were not predicted by this analysis.

**Figure 2.**
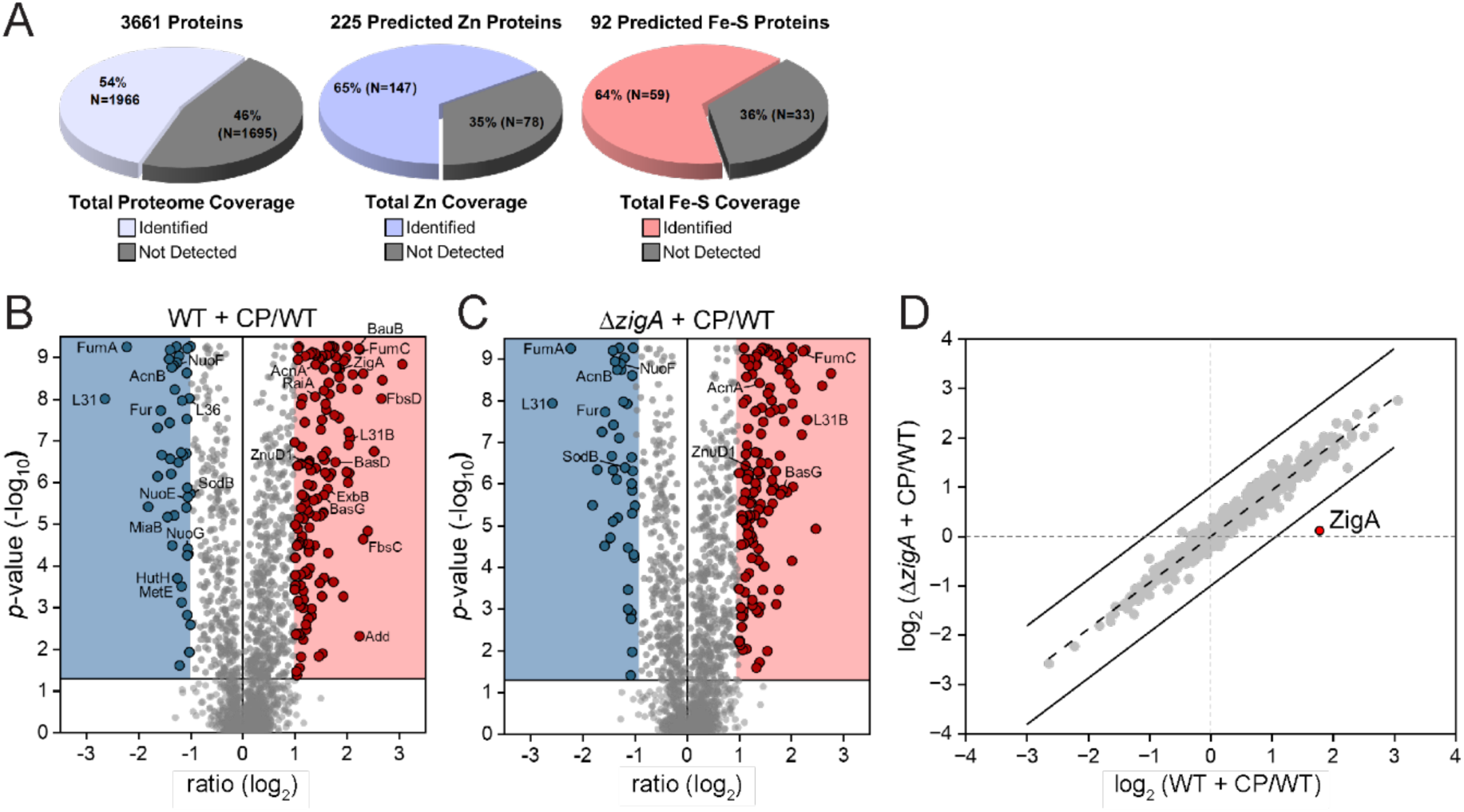
Change in protein abundance with CP treatment. (A) Pie charts that summarize the statistics of proteome and metalloproteome coverage in these experiments in *A. baumannii* lysates. (B) Volcano plot of WT cells treated with CP vs. untreated WT cells. (C) Volcano plot of Δ*zigA* cells treated with CP vs untreated WT cells. (D) log-log plot of CP-mediated changes in the proteomes of WT vs. Δ*zigA* cells. *Solid lines*, boundaries of a net fold change of 2 (log_2_ fold change = 1) deviating from the trendline (*dashed line*) with the ZigA data point indicated in red. In panel A, 141 proteins increase in cellular abundance ≥ 2-fold (*red* box) while 45 decrease in abundance ≥ 2-fold (*blue* box).

### Changes in Protein Abundance in CP-stressed *A. baumannii*

A volcano plot illustrating the impact of CP stress on the WT *A. baumanni* proteome shows major perturbations of the proteome. Of the 1966 total proteins detected, 141 proteins increased and 45 proteins decreased in abundance (Figure 2B, Document S2). The changes observed with the WT strain upon CP stress are virtually identical with those obtained in a pairwise comparison of the CP-stressed Δ*zigA* strain vs. untreated WT cells (Figure 2C-D). This reveals that the cellular impact of CP is not appreciably impacted by the absence of ZigA, which as a Zur target, reaches high intracellular levels (≈400 molecules per cell) under CP-stress (Figure S2; Table 1).^27,30^ Furthermore, ZigA expression is confirmed to be strongly repressed in unstressed cultures as indicated by a nearly unchanged protein abundance in the Δ*zigA*+CP/WT dataset (Document S2).

**Table 1.** List of selected proteins involved in Fe/Zn homeostasis or the stress response that show increased cell abundance in CP-stressed wild-type (WT+CP) or *ΔzigA (ΔzigA + CP)* vs. WT untreated control cells.^a^.

| Uniprot Entry | KEGG Entry | Description | Gene Name | Protein Ratio (log2) WT+CP v WT | Protein Ratio (log2) $\Delta zigA$+CP v WT | Fur/Zur<sup>b</sup> | Biological Function |
| --- | --- | --- | --- | --- | --- | --- | --- |
| A0A1E3MAV2 | A1S_2579 | 2,3-dihydro-2,3-dihydroxybenzoate dehydrogenase | *fbsD* | 2.66 | 2.20 | Fur | Fimsbactin biosynthesis |
| A0A1E3MAY3 | A1S_2570 | Acetyltransferase | *fbsK* | 2.52 | 2.03 | Fur | Fimsbactin biosynthesis |
| A0A1L5TK40 | A1S_2580 | 2,3-dihydro-2,3-dihydroxybenzoate synthetase | *fbsC* | 2.31 | 2.01 | Fur | Fimsbactin biosynthesis |
| A0A1E3M124 | A1S_0771 | Small-conductance mechanosensitive channel |  | 2.31 | 2.07 |  | Ion transport |
| V5VCJ3 | A1S_1986 | Fumarate hydratase class II | *fumC* | 2.24 | 2.22 |  | TCA cycle |
| A0A1E3M675 | A1S_2390 | Acinetobactin biosynthesis protein | *basB* | 2.23 | 2.27 | Fur | Acinetobactin biosynthesis |
| A0A059ZM74 | A1S_0391 | Large ribosomal subunit protein bL31B | *rpmE2* | 2.06 | 2.30 | Zur | L31 Zn-sparing paralog |
| A0A1E3MDH8 | A1S_2571 | Ornithine decarboxylase | *fbsJ* | 2.04 | 1.79 | Fur | Fimsbactin biosynthesis |
| A0A1E3MAZ6 | A1S_2572 | Alcaligin biosynthesis protein | *fbsI* | 2.03 | 1.63 | Fur | Fimsbactin biosynthesis |
| A0A5P1UDU4 | A1S_2581 | isochorismate synthase | *fbsB* | 2.03 | 1.71 | Fur | Fimsbactin biosynthesis |
| V5VE02 | A1S_1950 | Universal stress protein |  | 2.02 | 1.92 |  | Stress factor |
| A0A1E3M655 | A1S_2380 | Isochorismatase | *basF* | 2.01 | 2.01 | Fur | Acinetobactin biosynthesis |
| A0A5P1UDB1 | A1S_2574 | 2,3-dihydroxybenzoate-AMP ligase | *fbsH* | 1.99 | 1.64 | Fur | Fimsbactin biosynthesis |
| A0A1E3M5T3 | A1S_1412 | Glutathione S-transferase |  | 1.89 | 1.68 |  | ROS resistance |
| A0A059ZT15 | A1S_2072 | Universal stress protein |  | 1.85 | 2.02 |  | Stress factor |
| A0A1E3M686 | A1S_2384 | Histamine N-monooxygenase | *basC* | 1.79 | 1.76 | Fur | Acinetobactin biosynthesis |
| A0A1E3M8N0 | A1S_2381 | 2,3-dihydroxybenzoate-AMP ligase | *basE* | 1.78 | 1.78 | Fur | Acinetobactin biosynthesis |
| A0A7U4X7Y7 | A1S_3411 | CobW C-terminal domain-containing protein ZigA | *zigA* | 1.78 | 0.12 | Zur | Zn homeostasis |
| A0A1E3M6B6 | A1S_2383 | Peptide synthetase | *basD* | 1.78 | 1.70 | Fur | Acinetobactin biosynthesis |
| A0A1E3M653 | A1S_2386 | Ferric anguibactin-binding protein | *bauB* | 1.76 | 1.82 | Fur | Acinetobactin import |
| A0A1E3M3K8 | A1S_1654 | Oxaloacetate decarboxylase | *bfnG* | 1.76 | 1.55 | Fur | Baumannoferrin biosynthesis |
| V5VA41 | A1S_2863 | AhpC/TSA family protein | *ahpC* | 1.72 | 1.48 |  | ROS resistance |
| A0A5P1UFP4 | A1S_2566 | Ligand-gated channel protein | *fbsN* | 1.70 | 1.41 | Fur | Fimsbactin import |
| A0A1E3M3E1 | A1S_1652 | lucA/lucC family siderophore biosynthesis protein | *bfnE* | 1.67 | 1.44 | Fur | Baumannoferrin biosynthesis |
| A0A1E3M685 | A1S_2372 | Isochorismate synthase | *basJ* | 1.67 | 1.71 | Fur | Acinetobactin biosynthesis |
| V5VE85 | A1S_1470 | Glutathione peroxidase | *bsaA* | 1.66 | 1.62 |  | ROS resistance |
| A0A7U4X5T2 | A1S_2578 | Peptide synthetase | *fbsE* | 1.65 | 1.30 | Fur | Fimsbactin biosynthesis |
| A0A1E3M665 | A1S_2391 | Peptide synthetase | *basA* | 1.64 | 1.55 | Fur | Acinetobactin biosynthesis |
| V5VIG0 | A1S_0453 | Biopolymer transport protein ExbB | *exbB* | 1.63 | 1.66 | Zur | Zn uptake |
| V5VDG9 | A1S_1657 | Acetyltransferase | *bfnL* | 1.61 | 1.69 | Fur | Baumannoferrin biosynthesis |
| V5VEI3 | A1S_1246 | Universal stress family protein |  | 1.60 | 1.42 |  | Stress factor |
| A0A1E3M3G0 | A1S_1651 | Siderophore achromobactin biosynthesis protein AcsC | *bfnD* | 1.59 | 1.46 | Fur | Baumannoferrin biosynthesis |
| A0A7U4X5W9 | A1S_2575 | Peptide synthetase | *fbsG* | 1.58 | 1.21 | Fur | Fimsbactin biosynthesis |
| A0A1E3M7F7 | A1S_0558 | Aconitate hydratase | *acnA* | 1.56 | 1.39 |  | TCA cycle |
| A0A1E3M668 | A1S_2379 | Histidine decarboxylase | *basG* | 1.54 | 1.84 | Fur | Acinetobactin biosynthesis |
| A0A059ZIQ0 | A1S_1009 | Copper homeostasis protein | *nlpE* | 1.52 | 1.34 |  | ROS resistance |
| A0A1E3M3D7 | A1S_1648 | Ornithine monooxygenase | *bfnB* | 1.49 | 1.27 | Fur | Baumannoferrin biosynthesis |
| A0A1E3M8P2 | A1S_2392 | NADPH-dependent ferric siderophore reductase | *bauF* | 1.43 | 1.18 | Fur | Acinetobactin reductase |
| A0A1E3MBP6 | A1S_2892 | TonB-dependent receptor | *znuD1* | 1.40 | 1.38 |  | Zn uptake |
| V5VGU2 | A1S_0683 | Ribosome hibernation promoting factor | *raiA* | 1.39 | 1.31 |  | Ribosome hibernation |
| A0A1E3M7T8 | A1S_0452 | TonB-dependent receptor | *tonB* | 1.33 | 1.34 | Zur | Zn uptake |
| A0A5P1UI20 | A1S_2582 | AraC family transcriptional regulator | *fbsA* | 1.32 | 0.99 | Fur | Fimsbactin regulation |
| A0A1E3MCQ5 | A1S_2567 | Thioesterase | *ftsM* | 1.28 | 1.18 | Fur | Fimbractin biosynthesis |
| A0A1E3M630 | A1S_2378 | ABC transporter | *barA* | 1.26 | 1.37 | Fur | Acinetobactin export |
| A0A5P1UK94 | A1S_1647 | Siderophore biosynthesis protein | *bfnA* | 1.19 | 1.17 | Fur | Baumannoferrin biosynthesis |
| V5VGK1 | A1S_1186 | Chaperone protein ClpB | *clpB* | 1.15 | 1.14 |  | Stress factor |
| A0A059ZN38 | A1S_2387 | ATP-binding cassette domain-containing protein | *bauE* | 1.15 | 1.22 | Fur | Acinetobactin import |
| A0A1E3M643 | A1S_2374 | Thioesterase | *basH* | 1.14 | 1.36 | Fur | Acinetobactin biosynthesis |
| A0A1L5TMI2 | A1S_0146 | Zinc ABC transporter substrate-binding protein | *znuA* | 1.14 | 1.10 |  | Zn uptake |
| V5V9K7 | A1S_2834 | Large-conductance mechanosensitive channel |  | 1.12 | 1.05 |  | Ion transport |
| V5VH78 | A1S_0454 | Biopolymer transport ExbD/TolR family protein | *exbD* | 1.11 | 1.12 | Zur | Zn uptake |
| A0A0E1FM87 | A1S_0474 | Ferric siderophore receptor protein | *bfrD* | 1.11 | 0.99 | Fur | Siderophore uptake |
| V5VI41 | A1S_0034 | 2,3-dihydroxybenzoate-2,3-dehydrogenase | *yciK* | 1.08 | 1.20 | Fur |  |
| A0A1E3M7N8 | A1S_0412 | Catalase-peroxidase | *katG* | 1.06 | 0.97 |  | Stress factor |
| V5VHL8 | A1S_0136 | Glutathione-S-transferase | *dcmA* | 1.06 | 0.92 |  | Stress factor |
| A0A1E3M8A1 | A1S_0980 | Outer membrane receptor protein | *pirA* | 1.06 | 1.10 |  | Siderophore uptake |
| A0A1E3MBF3 | A1S_2765 | Glyoxalase III (protein deglycase) | *hchA* | 1.05 | 1.13 |  | Stress factor |
| A0A7U4XBR3 | A1S_2076 | Outer membrane porin |  | 1.04 | 0.99 |  | Fe(III)-coprogen receptor |
| V5VA09 | A1S_2692 | Universal stress protein |  | 1.01 | 1.08 |  | Stress factor |
| A0A1E3M2Y5 | A1S_1386 | Catalase | *katE* | 1.00 | 0.83 |  | ROS resistance |
| A0A1E3M5V4 | A1S_1411 | Glutathione-dependent disulfide-bond oxidoreductase |  | 1.00 | 0.88 |  | Stress factor |

<sup>a</sup>See Document S2 for a list of all proteins
<sup>b</sup>Genes known to be regulated by ferric uptake repressor (Fur) or zinc uptake repressor (Zur).

Examination of the proteins that display increased abundance under CP stress is consistent with the pronounced Zn- and Fe-starvation phenotype reported previously,^30^ as evidenced by the large number of proteins (26% of the total) that are components of the ferric uptake repressor (Fur) and Zur regulons (Table 1; Figure 3A). For example, the Fur-regulated enzymes used to synthesize the major mixed imidazole-catecholate Fe(III) siderophore acinetobactin (Figure S3A) (BasA-J), and the inner membrane transporter required to uptake Fe(III)-acinetobactin complexes (BauA-E),^64–65^ as well as the Fe(III)-acinetobactin ferric reductase (BauF),^66^ all become significantly more abundant in CP-stressed cells (Figure S3B). In addition, a TonB-dependent outer membrane-dependent receptor and ExbB/ExbD, which collectively energize the outer membrane for ferric-siderophore transport, are also more abundant under these conditions (Table 1; Figure S3). Proteins and enzymes required for the biosynthesis and trafficking of the minor mixed catechol-hydroxymate Fe(III) siderophores, fimsbactin A and B (FbsA-E, FbsH-K, FbsM) as well as the baumannoferrin A and B, also become more abundant (Table 1; Figure S3). The Zur regulon is also highly induced by CP, and includes known Zur target genes encoding the low-Zn ribosomal protein paralog L31B (*rpmE2*), ZigA,^27^ and components of the high affinity zinc uptake system, including ZnuA, ZnuD1 and the TonB/ExbB/ExbD system that is regulated by Zur (Table 1).^67–68^

**Figure 3.**
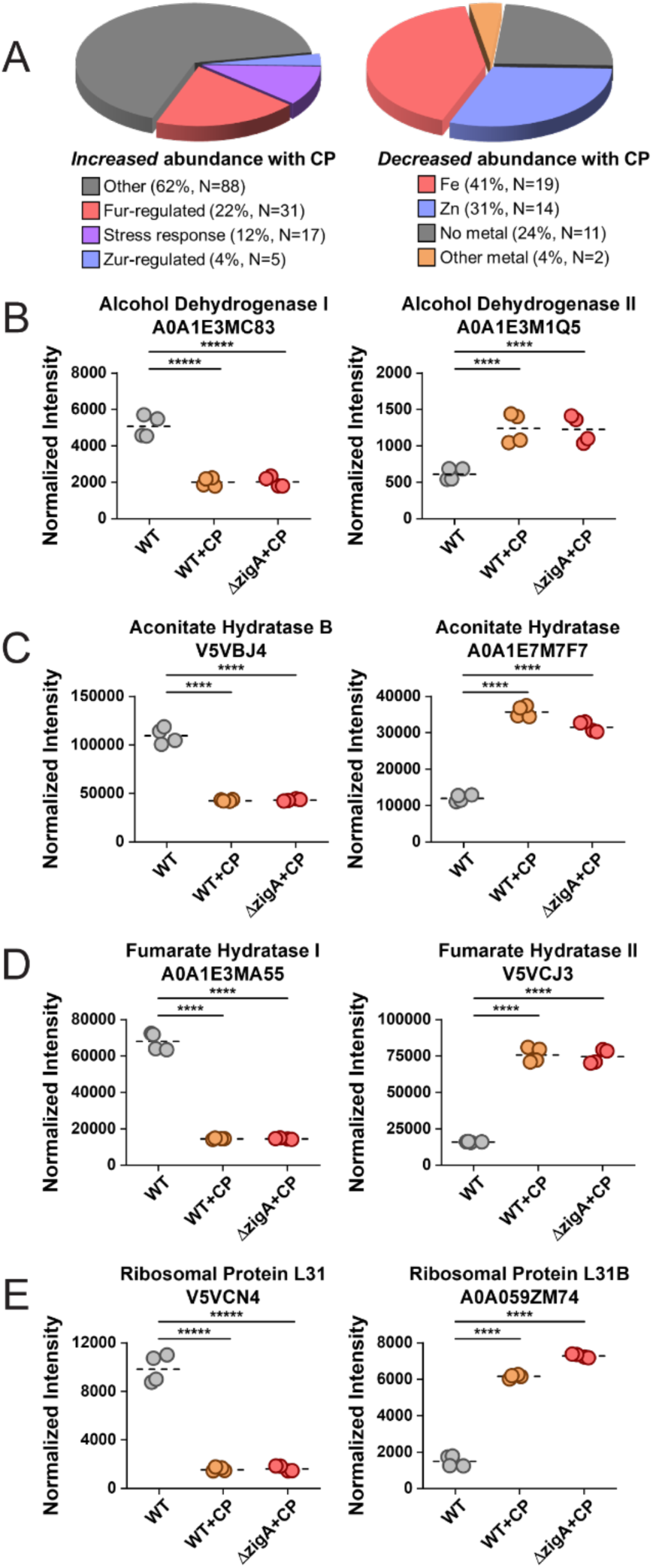
Changes in protein abundance upon CP treatment of WT and Δ*zigA* cells. (A) Pie charts that summarize the statistical and functional characteristics of *A. baumannii* proteins that increase (*left*; 141 proteins total) or decrease (*right*; 45 proteins) in cell abundance in CP stressed WT cells. (B)-(E) Normalized intensity of selected protein paralogs that exhibit reciprocal regulation in WT (housekeeping) vs. CP-stressed cells. Trivial names for each protein and their Uniprot numbers are indicated. Statistical significance according to an unpaired *t*-test is indicated as follows:*, *p*≤0.05; **, *p*≤0.01; ***, *p*≤0.001; ****, *p*≤0.0001

Closer examination of the 45 proteins that display significantly decreased abundance under conditions of CP stress reveals a high enrichment (76% of the total) of known metal (Zn, Fe) or metallocofactor (Fe-S cluster) binding proteins and proteins involved in Fe-S protein biogenesis, including the cysteine desulfurase IscS, and the Fe-S assembly scaffold protein IscU (Table 2; Figure 3A). Fur itself is decreased by ≈3-fold, which is consistent with increased expression of the Fur regulon under these conditions (Table 1; Figure 2B). Proteins with decreased abundance include the large ribosomal protein L31 (encoded by *rpmE*), and a number of cell abundant, cytoplasmic [4Fe-4S] proteins (see Discussion), including the citric acid cycle housekeeping enzymes aconitase B (AcnB) and the class I fumarase (FumA), as well as a functionally uncharacterized, canonical Zn-dependent alcohol dehydrogenase. In fact, all four of these proteins are reciprocally regulated, in which a companion “low-metal paralog”^51^ encoded by another gene displays increased abundance under conditions of CP-stress (Figure 3B-E).

**Table 2.**
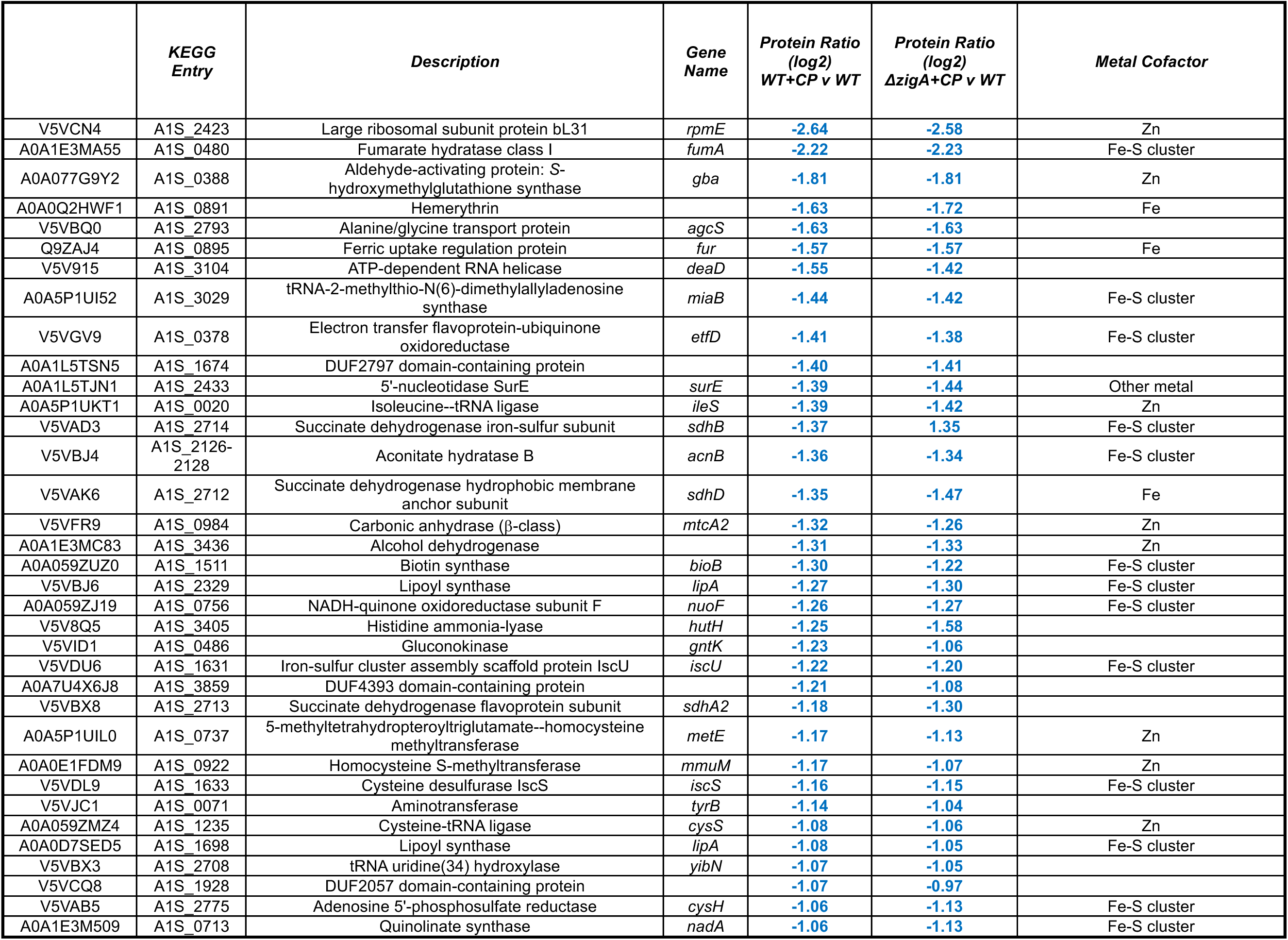
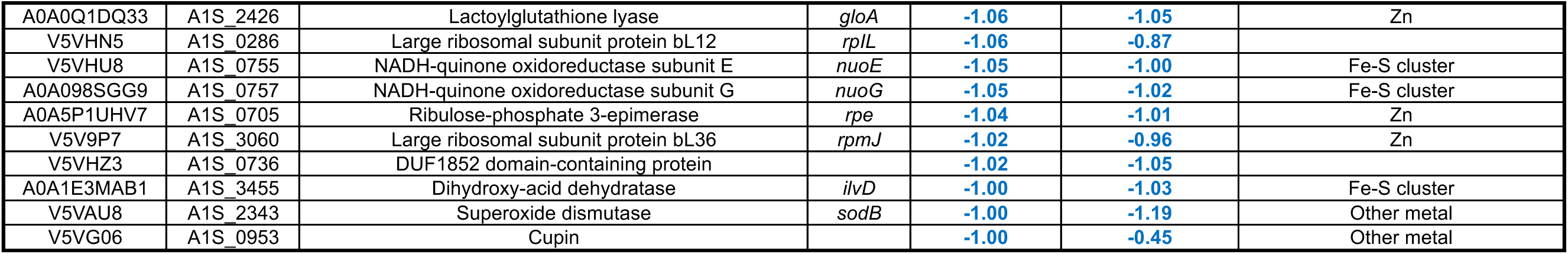
List of all proteins that show decreased abundance in CP-stressed wild-type (WT+CP) or *ΔzigA (ΔzigA + CP)* vs. WT untreated control cells.

|  | **KEGG**<br>**Entry** | **Description** | **Gene**<br>**Name** | **Protein Ratio**<br>**(log2)**<br>**WT+CP v WT** | **Protein Ratio**<br>**(log2)**<br>**ΔzigA+CP v WT** | **Metal Cofactor** |
| --- | --- | --- | --- | --- | --- | --- |
| V5VCN4 | A1S_2423 | Large ribosomal subunit protein bL31 | *rpmE* | -2.64 | -2.58 | Zn |
| A0A1E3MA55 | A1S_0480 | Fumarate hydratase class I | *fumA* | -2.22 | -2.23 | Fe-S cluster |
| A0A077G9Y2 | A1S_0388 | Aldehyde-activating protein: S-hydroxymethylglutathione synthase | *gba* | -1.81 | -1.81 | Zn |
| A0A0Q2HWF1 | A1S_0891 | Hemerythrin |  | -1.63 | -1.72 | Fe |
| V5VBQ0 | A1S_2793 | Alanine/glycine transport protein | *agcS* | -1.63 | -1.63 |  |
| Q9ZAJ4 | A1S_0895 | Ferric uptake regulation protein | *fur* | -1.57 | -1.57 | Fe |
| V5V915 | A1S_3104 | ATP-dependent RNA helicase | *deaD* | -1.55 | -1.42 |  |
| A0A5P1UI52 | A1S_3029 | tRNA-2-methylthio-N(6)-dimethylallyl adenosine synthase | *miaB* | -1.44 | -1.42 | Fe-S cluster |
| V5VGV9 | A1S_0378 | Electron transfer flavoprotein-ubiquinone oxidoreductase | *etfD* | -1.41 | -1.38 | Fe-S cluster |
| A0A1L5TSN5 | A1S_1674 | DUF2797 domain-containing protein |  | -1.40 | -1.41 |  |
| A0A1L5TJN1 | A1S_2433 | 5'-nucleotidase SurE | *surE* | -1.39 | -1.44 | Other metal |
| A0A5P1UKT1 | A1S_0020 | Isoleucine--tRNA ligase | *ileS* | -1.39 | -1.42 | Zn |
| V5VAD3 | A1S_2714 | Succinate dehydrogenase iron-sulfur subunit | *sdhB* | -1.37 | 1.35 | Fe-S cluster |
| V5VBJ4 | A1S_2126-2128 | Aconitate hydratase B | *acnB* | -1.36 | -1.34 | Fe-S cluster |
| V5VAK6 | A1S_2712 | Succinate dehydrogenase hydrophobic membrane anchor subunit | *sdhD* | -1.35 | -1.47 | Fe |
| V5VFR9 | A1S_0984 | Carbonic anhydrase (β-class) | *mtcA2* | -1.32 | -1.26 | Zn |
| A0A1E3MC83 | A1S_3436 | Alcohol dehydrogenase |  | -1.31 | -1.33 | Zn |
| A0A059ZUZ0 | A1S_1511 | Biotin synthase | *bioB* | -1.30 | -1.22 | Fe-S cluster |
| V5VBJ6 | A1S_2329 | Lipoyl synthase | *lipA* | -1.27 | -1.30 | Fe-S cluster |
| A0A059ZJ19 | A1S_0756 | NADH-quinone oxidoreductase subunit F | *nuoF* | -1.26 | -1.27 | Fe-S cluster |
| V5V8Q5 | A1S_3405 | Histidine ammonia-lyase | *hutH* | -1.25 | -1.58 |  |
| V5VID1 | A1S_0486 | Gluconokinase | *gntK* | -1.23 | -1.06 |  |
| V5VDU6 | A1S_1631 | Iron-sulfur cluster assembly scaffold protein IscU | *iscU* | -1.22 | -1.20 | Fe-S cluster |
| A0A7U4X6J8 | A1S_3859 | DUF4393 domain-containing protein |  | -1.21 | -1.08 |  |
| V5VBX8 | A1S_2713 | Succinate dehydrogenase flavoprotein subunit | *sdhA2* | -1.18 | -1.30 |  |
| A0A5P1UII0 | A1S_0737 | 5-methyltetrahydropteroyltrimethylglutamate--homocysteine methyltransferase | *metE* | -1.17 | -1.13 | Zn |
| A0A0E1FDM9 | A1S_0922 | Homocysteine S-methyltransferase | *mmuM* | -1.17 | -1.07 | Zn |
| V5VDL9 | A1S_1633 | Cysteine desulfurase IscS | *iscS* | -1.16 | -1.15 | Fe-S cluster |
| V5VJC1 | A1S_0071 | Aminotransferase | *tyrB* | -1.14 | -1.04 |  |
| A0A059ZMZ4 | A1S_1235 | Cysteine-tRNA ligase | *cysS* | -1.08 | -1.06 | Zn |
| A0A0D7SED5 | A1S_1698 | Lipoyl synthase | *lipA* | -1.08 | -1.05 | Fe-S cluster |
| V5VBX3 | A1S_2708 | tRNA uridine(34) hydroxylase | *yibN* | -1.07 | -1.05 |  |
| V5VCQ8 | A1S_1928 | DUF2057 domain-containing protein |  | -1.07 | -0.97 |  |
| V5VAB5 | A1S_2775 | Adenosine 5'-phosphosulfate reductase | *cysH* | -1.06 | -1.13 | Fe-S cluster |
| A0A1E3M509 | A1S_0713 | Quinolinate synthase | *nadA* | -1.06 | -1.13 | Fe-S cluster |
| A0A0Q1DQ33 | A1S_2426 | Lactoylglutathione lyase | *gloA* | -1.06 | -1.05 | Zn |
| V5VHN5 | A1S_0286 | Large ribosomal subunit protein bL12 | *rplL* | -1.06 | -0.87 |  |
| V5VHU8 | A1S_0755 | NADH-quinone oxidoreductase subunit E | *nuoE* | -1.05 | -1.00 | Fe-S cluster |
| A0A098SGG9 | A1S_0757 | NADH-quinone oxidoreductase subunit G | *nuoG* | -1.05 | -1.02 | Fe-S cluster |
| A0A5P1UHV7 | A1S_0705 | Ribulose-phosphate 3-epimerase | *rpe* | -1.04 | -1.01 | Zn |
| V5V9P7 | A1S_3060 | Large ribosomal subunit protein bL36 | *rpmJ* | -1.02 | -0.96 | Zn |
| V5VHZ3 | A1S_0736 | DUF1852 domain-containing protein |  | -1.02 | -1.05 |  |
| A0A1E3MAB1 | A1S_3455 | Dihydroxy-acid dehydratase | *ilvD* | -1.00 | -1.03 | Fe-S cluster |
| V5VAU8 | A1S_2343 | Superoxide dismutase | *sodB* | -1.00 | -1.19 | Other metal |
| V5VG06 | A1S_0953 | Cupin |  | -1.00 | -0.45 | Other metal |

Biochemical experiments validate this aconitase switch as we find that low-metal induced paralog AcnA activity is completely refractory to chelator (EDTA) challenge in contrast to the housekeeping enzyme, AcnB (Figure 4A; Figure S4); these findings mirror what was found in *E. coli*, which in that work was traced to a weaker Fe-binding affinity.^69^ The enhanced expression of a “back-up” aconitase A (*acnA*) has not been previously reported in *A. baumannii*, and could be regulated post-transcriptionally, as recently described in *Staphylococcus aureus*.^50^ The reciprocal transcriptional regulation of the class I fumarase and Fe-independent class II fumarase, FumC, was previously noted under conditions of iron starvation in *A. baumannii* ATCC 17978,^70^ as has the reciprocal transcription of L31 vs the Zur-regulated low-Zn paralog L31B (encoded by *rpmE2*).^30^ In addition to these enzymes, two cofactor-synthesizing [4Fe-4S] enzymes, biotin synthase and lipoyl synthase, and a Zn-dependent carbonic anhydrase (MtcA2) also decrease in abundance (Table 2). Two other Zn-requiring methyltransferases involved in methionine synthesis (MetE) or homocysteine catabolism (MmnE) become less abundant upon CP-stress, as does the Zn-utilizing enzyme histidine ammonia lyase (HutH), previously tied to the CP-stress response (Table 2).^27^

**Figure 4.**
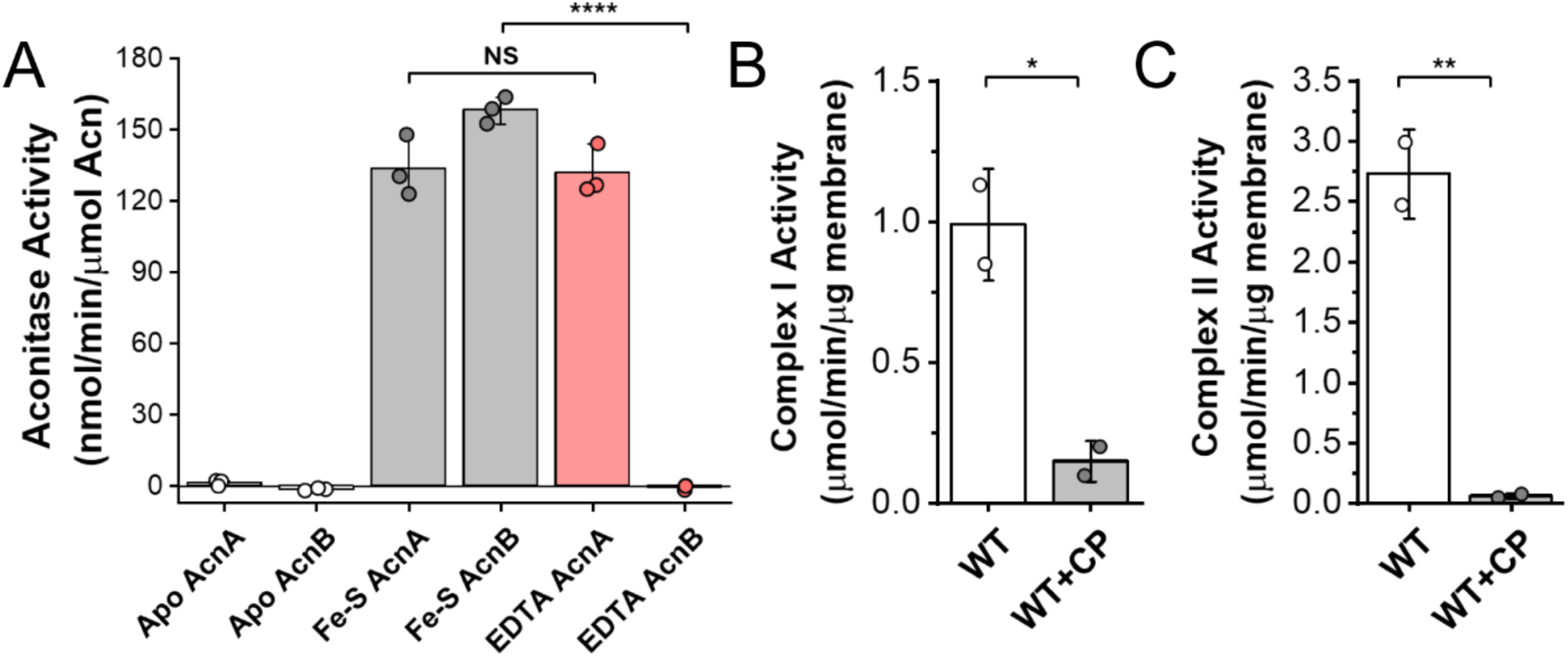
Measured activities of *A. baumannii* aconitases, complex I, and complex II. (A) Enzyme kinetics of AcnA and AcnB that have been chemically reconstituted with Fe-S clusters and treated with EDTA. (B) total membrane NADH:ubiquinone dehydrogenase (complex I) activities in biological duplicate as measured by reduction of NADH at 340 nm in untreated and (WT) an CP-stressed cells (WT+CP). (C) Total membrane succinate dehydrogenase:ubiquinone oxidoreductase (complex II) activities as measured by reduction of succinate-2,6-dichlorophenolindophenol (DCPIP) at 600 nm in biological duplicate. Significance calculated from an unpaired *t*-test, *, *p*≤0.05; **, *p*≤0.01; ***, *p*≤0.001; ****, *p*≤0.0001.

CP-stressed cells also show significant reduction in Fe-S-cluster-containing membrane proteins including NuoE, NuoF and NuoG, key components of respiratory complex I, the NADH-ubiquinone oxidoreductase, which allows NADH-carried electrons to enter the electron transfer chain (ETC) (Table 2). In addition, comparable decreases in abundance are observed for respiratory complex II subunits that make up succinate dehydrogenase-ubiquinone oxidoreductase, the ETC entry point for FADH_2_-derived electrons (Table 2). These findings again mirror results obtained from *E. coli*, where Fe-starvation leads to decreased abundance of components of the respiratory complexes I and II.^71^ Both activities are significantly decreased in CP-stressed cells (Figure 4B-C; Figure S5) (see below). In summary, we conclude that at least some fraction of the metalloproteome is destabilized and thus subject to increased turnover. This might be exacerbated by undermetalation of the essential Zn site II in the protein chaperone DnaJ, which would reduce the ability of cells to rescue misfolded proteins (discussed below).

### Changes in Cysteine Reactivity Across the Proteome

We next determined the abundance-corrected changes in Cys reactivity across the proteome (Figure 5; Table 3; Table S2). Of the 2645 total unique quantified Cys-containing peptides, 150 unique Cys increase and 67 unique Cys decrease in net reactivity (≥2-fold; χ_2_≥9) in WT *A. baumannii* stressed with CP (Figure 5A; Document S2). Once again, it is difficult to broadly differentiate the CP-treated WT from CP-treated Δ*zigA* cells (Figure 5B-C), with the changes in Cys reactivity very nearly identical in the two strain backgrounds (Table 3). In no case did we see a large increase in reactivity of a metal-coordinating Cys only in the Δ*zigA* strain, as might be expected for a ZigA metallochaperone client.^3,72^ This suggests that if a high abundance client exists for ZigA, it likely involves a metal-coordination site that lacks Cys ligation, or the relevant Cys-containing peptide was not detectable in these experiments.

**Figure 5.**
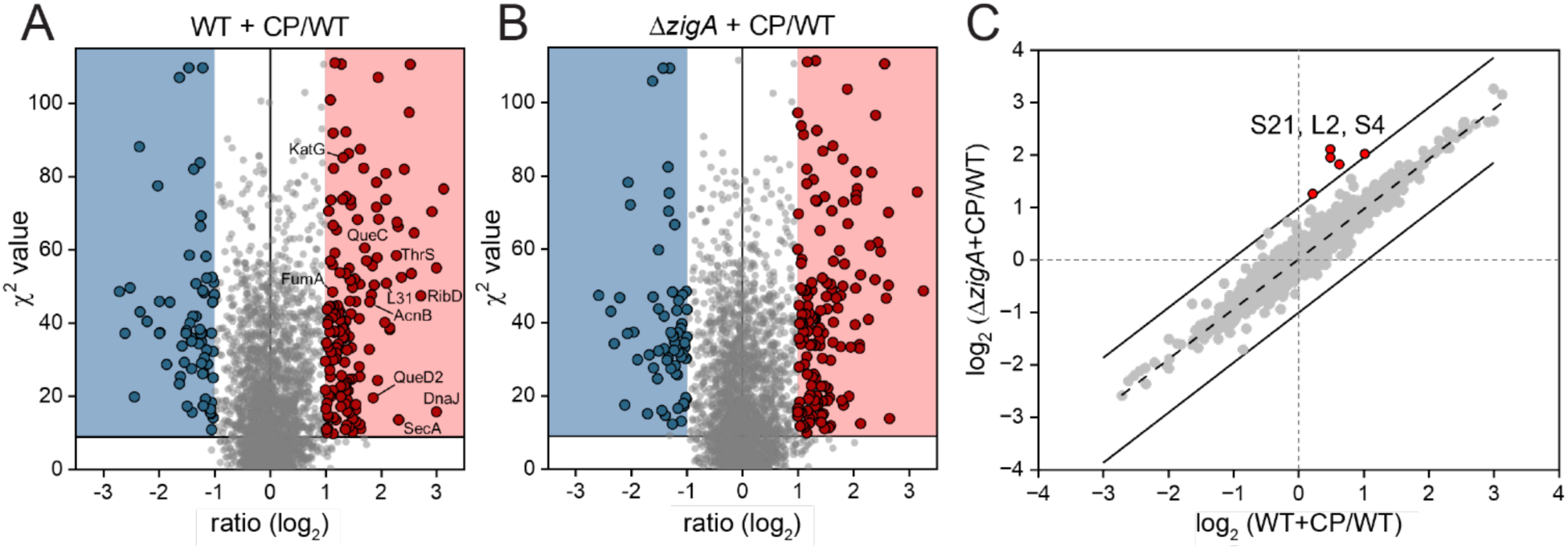
Change in net Cys reactivity in the proteome with CP treatment. A) wild-type cells treated with CP vs. untreated WT cells; (B) Δ*zigA* cells treated with CP vs. untreated WT cells. (C) log-log plot of CP-mediated changes in Cys reactivity in the proteomes of WT vs. Δ*zigA* cells. *Solid lines*, boundaries of a net fold change of 2 (log_2_ fold change = 1) deviating from the trendline (*dashed line*) with the five peptides, corresponding to just three unique Cys, just outside the range indicated; all belong to three ribosomal proteins.

**Table 3.** List of Zn or Fe-S cluster harboring proteins and their Cys-coordinating residues that show increased reactivity in CP-stressed wild-type (WT+CP) or CP-stressed Δ*zigA* (Δ*zigA* + CP) vs. WT untreated control cells^a^.

| *Uniprot Entry* | *KEGG Entry* | *Description* | *Gene Name* | *Modification Position (Cys)* | *Net Cys Ratio (log2) WT+CP v WT* | *Net Cys Ratio (log2) $\Delta zigA$+CP v WT* | *Metal Cofactor* |
| --- | --- | --- | --- | --- | --- | --- | --- |
| V5V8K0 | A1S_3443 | Chaperone protein DnaJ | *dnaJ* | 164 | **3.00** | **2.65** | Zn |
| A0A5P1UGY3 | A1S_0221 | Riboflavin biosynthesis protein RibD | *ribD* | 88 | **2.72** | **2.59** | Zn |
| A0A059ZTZ6 | A1S_0592 | Threonine--tRNA ligase | *thrRS* | 333\|335 | **2.55** | **2.63** | Zn |
| A0A0H4UCN7 | A1S_3619 | Zinc ribbon domain-containing protein |  | 6 | **2.16** | **2.11** | Zn |
| A0A1E3MBL8 | A1S_2821 | Alkylphosphonate utilization protein |  | 6 | **2.10** | **2.04** | Zn |
| A0A1E3MBS1 | A1S_2862 | Protein translocase subunit SecA | *secA* | 903 | **2.32** | **2.13** | Zn |
| A0A086HW53 | A1S_0531 | Small ribosomal subunit biogenesis GTPase RsgA | *engC* | 319 | **2.07** | **2.03** | Zn |
| A0A1V3DEG5 | A1S_2542 | 7-cyano-7-deazaguanine synthase | *queC* | 190 | **1.93** | **1.84** | Zn |
| V5VCN4 | A1S_2423 | Large ribosomal subunit protein bL31 | *rpmE* | 38\|41 | **1.88** | **1.81** | Zn |
| A0A081GYS3 | A1S_1566 | 6-carboxy-5,6,7,8-tetrahydropterin synthase | *queD2* | 18 | **1.86** | **1.54** | Zn |
| V5VBJ4 | A1S_2126-2128 | Aconitate hydratase B | *acnB* | 718 | **1.80** | **1.72** | Fe-S cluster |
| V5VAD3 | A1S_2714 | Succinate dehydrogenase iron-sulfur subunit | *sdhB* | 210\|214 | **1.79** | **1.77** | Fe-S cluster |
| A0A059ZTE3 | A1S_1176 | Alanine--tRNA ligase | *alaRS* | 670 | **1.71** | **1.75** | Zn |
| A0A0H4UDV2 | A1S_2114 | DNA mismatch repair protein MutL | *mutL* | 596 | **1.49** | **1.42** | Zn |
| A0A059ZMZ4 | A1S_1235 | Cysteine--tRNA ligase | *cysRS* | 30 | **1.47** | **1.63** | Zn |
| A0A059ZUZ0 | A1S_1511 | Biotin synthase | *bioB* | 128 | **1.46** | **1.41** | Fe-S cluster |
| A0A077G9Y2 | A1S_0388 | S-hydroxymethylglutathione synthase | *gfa* | 33 | **1.39** | **1.59** | Zn |
| V5VFR9 | A1S_0984 | Carbonic anhydrase (β-class) | *mtcA2* | 50 | **1.37** | **1.19** | Zn |
| A0A1E3M178 | A1S_0778 | Methionine--tRNA ligase | *metRS* | 142 | **1.37** | **1.25** | Zn |
| V5VGB8 | A1S_0696 | ADP-ribose pyrophosphatase |  | 26\|29 | **1.29** | **1.32** | Zn |
| V5V9P7 | A1S_3060 | Large ribosomal subunit protein bL36 | *rpmJ* | 11 | **1.26** | **1.35** | Zn |
| A0A1E3M9N2 | A1S_0403 | Ribonuclease E | *rne* | 401 | **1.19** | **1.12** | Zn |
| A0A098SGG9 | A1S_0757 | NADH-quinone oxidoreductase | *nuoG* | 235 | **1.13** | **1.26** | Fe-S cluster |
| A0A1E3MA55 | A1S_0480 | Fumarate hydratase class I | *fumA* | 264\|277 | **1.13** | **1.07** | Fe-S cluster |
| V5VBH1 | A1S_2366 | GTP cyclohydrolase 1 | *folE* | 144 | **1.09** | **1.24** | Zn |
| V5VBJ6 | A1S_2329 | Lipoyl synthase | *lipA* | 95\|98 | **1.07** | **0.98** | Fe-S cluster |
| V5VI79 | A1S_0579 | GTP pyrophosphokinase | *relA* | 662 | **1.05** | **1.08** | Zn |
| A0A5P1UI52 | A1S_3029 | tRNA-2-methylthio-N(6)-dimethylallyladenosine synthase | *miaB* | 40 | **1.05** | **1.29** | Fe-S cluster |
| A0A5P1UKT1 | A1S_0020 | Isoleucine--tRNA ligase | *ileRS* | 418 | **1.03** | **1.02** | Zn |

<sup>a</sup>The Cys shown corresponds to the highest net change in Cys reactivity in the chelate. If two Cys are shown, these correspond to Cys within the same tryptic peptide. The net reactivity changes of other Cys residues associated with these chelates are given in Table S2.

We next carried out a statistical analysis to determine the fraction of the Zn and Fe-S-cluster proteome being probed in this experiment (Figure 6). Figure 6A shows that of the 106 Zn metalloproteins detected in this experiment, 31 harbor a Cys in the first coordination shell (Figure 6A, Table S2). Of these 31 proteins, 22 (76%) display a two-fold or greater increase in reactivity with a chi-squared greater than 9, indicative of loss of the metal cofactor (Figure 6A). To probe the relationship between relative protein abundance and undermetalation, we performed label-free proteomics on soluble cell lysates of unstressed WT, ranked proteins according to their fractional intensities^73–74^ (Document S3) and highlighted those that were undermetalated in our chemoproteomics datasets (Figure 7A). These 22 Zn proteins in CP-treated cells are present at a median cell abundance that is comparable to all detected proteins and detected Zn enzymes (Figures 7B-C). Interestingly, the median abundance of these 22 undermetalated enzymes is slightly lower than from the seven Zn enzymes that show no increase in fractional occupancy of the metal site (Figure 7D). These data show that it is not uniquely the most abundant Zn enzymes in unstressed cells that become undermetalated in CP-stressed cells.

**Figure 6.**
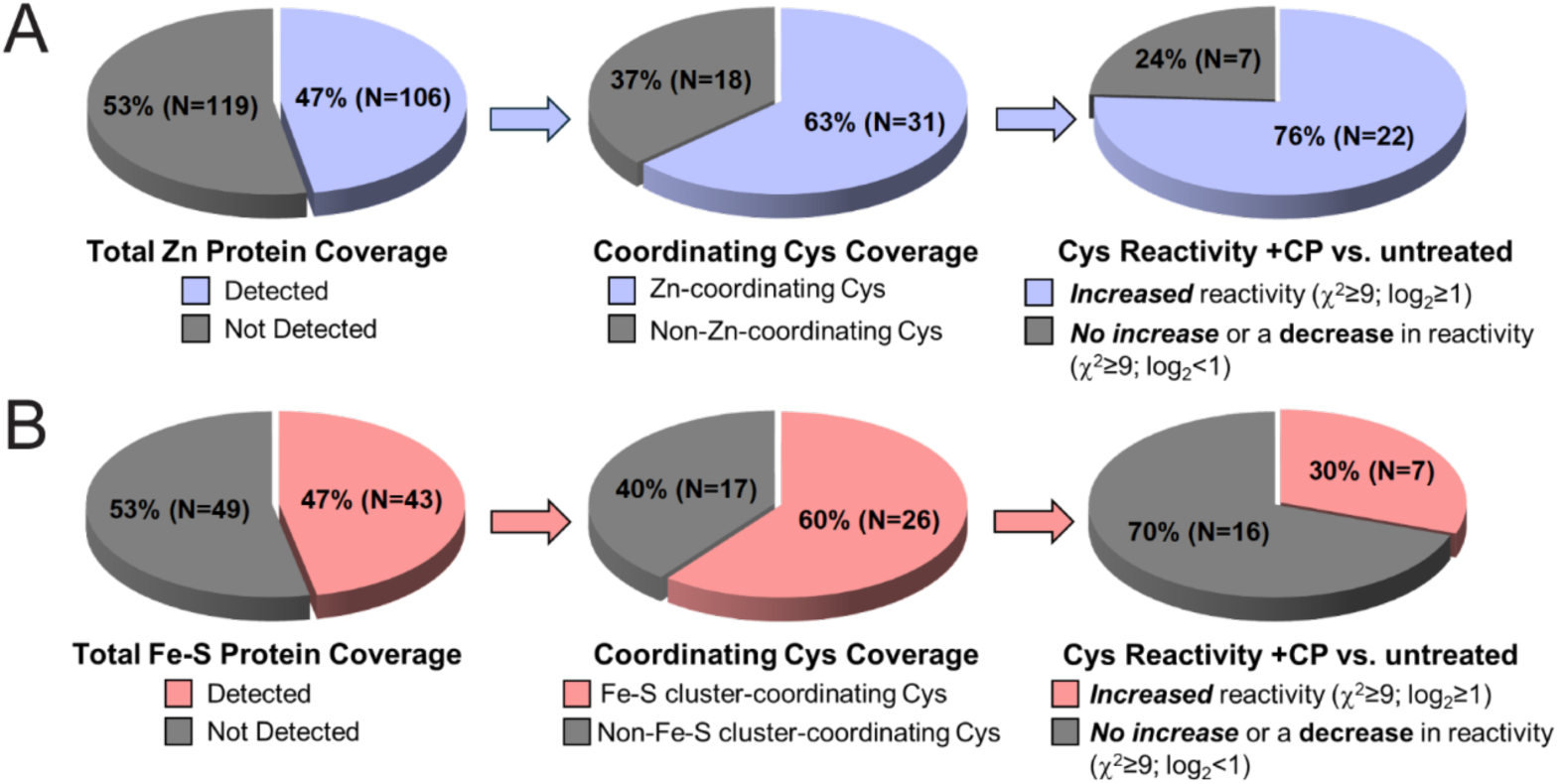
Statistical analysis of the quantifiable changes in Cys reactivity across the proteome comparing CP-treated vs. untreated WT cells. (A) Analysis of the Zn metalloproteome. Of the 225 Zn proteins (Document S1), 106 were detected at the peptide level in the Cys reactome (*left* pie chart). 49 of these Zn proteins are predicted to contain a Cys in the first coordination shell, and 31 Cys-containing peptides corresponded to a Zn-ligated Cys (*middle*). 22 out of 29 Cys exhibited a statistically significant change in Cys reactivity using the cut-offs indicated (*right*). An additional four Zn enzymes are identified in this analysis if log_2_ cut-off is dropped to 0.64 (1.5-fold increase in Cys reactivity) (Table S1). (B) Analysis of the Fe-S cluster proteome usijng the same filtering strategy. Of the 92 predicted Fe-S cluster proteins (Document S1), 43 are detected at the peptide level (*left*). 26 of these 43 proteins had a peptide that corresponded to an Fe-S cluster-ligated Cys (*middle*). 7 out of these 23 Cys show a statistically significant change in Cys reactivity using the cut-offs indicated (*right*). An additional four Fe-S proteins are identified in this analysis if log_2_ cut-off is dropped to 0.64 (1.5-fold increase in Cys reactivity) (Table S1).

**Figure 7.**
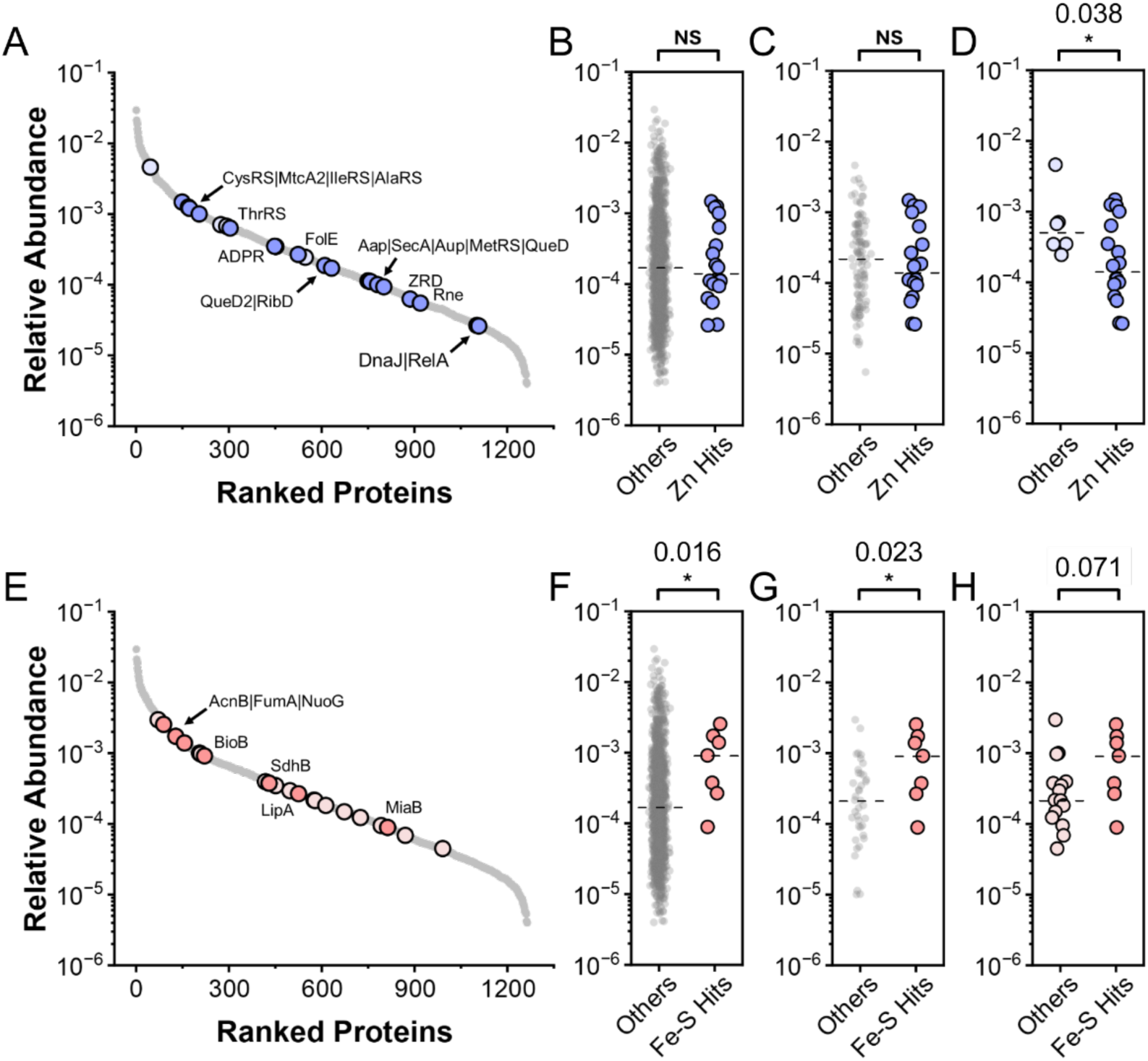
Analysis of the relative abundance of Zn (A)-(D) and Fe-S cluster (E)-(H) proteins for which Cys reactivity in the first coordination shell could be quantified [panels (A) and (E)] against all proteins in the proteome [panels (B) and (F)], all other Zn or Fe-S metalloproteins detected [panels (C) and (G), respetively], or other metal-sites in Zn- or Fe-S proteins where cysteine reactivity did not reach our significance threshold [panels (D) and (H), respectively]. Filled, dark circles, those metalloproteins that exhibit decreased metal occupancy; light filled circles, those that exhibit no change or decreased Cys reactivity with CP stress. Dotted lines represent the median abundance of each group, with the significance of difference in the medians calculated from a Mann-Whitney *U* analysis and the *p*-values listed above each comparison, *, *p*≤0.05; NS, not significant.

The opposite behavior seems to characterize the Fe-S proteome (Figure 7E-H). Out of the 43 Fe-S cluster proteins detected, 26 (60%) had a corresponding Cys-containing peptides that was localized to the first Fe-S cofactor coordination shell (Figure 6B). Only seven of these proteins are undermetalated by more than two-fold (Figure 6B), and these enzymes appear higher in median cell abundance than those Fe-S cluster coordination sites that do not reach this significance threshold for a change in reactivity in the cell (Figure 7E-G). This is in direct contrast with that of the Zn metalloproteome (Fig. 7A-C). Further, all seven enzymes that are severely undermetalated are present at lower levels in chronically CP-stressed cells which suggests that the Fe-S proteome might be subject to turnover when not fully metalated (Table 2), *i.e*., a global Fe-sparing response.^75^ This is not true of the undermetalated Zn proteome, with just four of the 22 on this list of “destabilized” Zn metalloproteins including L31, whch may constitute the bulk of the Zn-mobilization response.^76^ Finally, the median cell abundance of these seven undermetalated Fe-S cluster enzymes in unstressed *A. baumannii* trends toward higher levels relative to those proteins that show no increase in fractional occupancy of the metal site (Figure 7H); this is also in contrast to the Zn proteome (Figure 7D). This analysis suggests the interesting finding that the response of the Zn proteome to decreased cell-associated Zn is distinct from that of the behaavior of the Fe-S cluster proteome, which reflects no difference in total cell-associated Fe (Figure S1B).

Just below the 2-fold increase in Cys reactivity cut-off, are several other metalloproteins of interest (Table S1). These include IscU (C37), the only known Fe-S cluster assembly scaffold protein (log_2_=0.64) which becomes ≈2-fold more reactive in Δ*zigA* cells; IscU is also lower in cellular abundance (Table 2). In addition, the cysteine desulfurase IscS that functions upstream of IscU^77^ is also reduced in cell abundance under conditions of CP-stress (Table 2). These findings suggest a negative impact on Fe-S biogenesis under these conditions. In addition, a 4Fe-4S dicluster domain-containing (ferredoxin-like) enzyme annotated as RnfB of unknown function is also substantially undermetalated under these conditions (log_2_=0.94). This is in contrast to two other ferredoxins, WrbA and FdxB, that are not substantially impacted by CP stress (Document S2). Another site of interest is C92 (log_2_=0.93) of peptide deformylase (PDF), an essential mononuclear His_2_-Cys-H_2_O Fe(II) enzyme^78^ that co-translationally cleaves the *N*-formyl group of the initiator formyl-Met of the nascent chain, the first step in the N-terminal Met excision pathway. PDF remains a *bona fide* antimicrobial drug target in many bacteria, including recent work in *A. baumannii,*^79^ and oxidation or over-oxidation of the Fe(II)-coordinating Cys leads to inactivation of the enzyme in *Salmonella enterica*.^78,80^

### Undermetalation of the Fe-S Proteome, Cofactor Biosynthesis and Energy Generation

One subset of enzymes not previously discussed that is affected by CP-mediated Fe starvation is the abundant Fe-S cluster proteins associated with a) biosynthesis of cofactors utilized by the energy-generating TCA cycle enzymes (biotin ligase, lipoate synthase), b) enzymes of the TCA cycle itself (aconitase, fumarase and succinate dehydrogenase), c) enzymes for CO_2_ fixation (MtcA2), and d) subunits of respiratory complexes I and II, which utilize FADH_2_ and NADH, respectively, as a source of electrons to power the membrane potential and ATP synthesis (Figure 8). We find significantly reduced specific activities for both respiratory complexes I and II (Figure 4B-C; Figure S5), derived from reduced protein levels, and a significantly reduced metal content in what actually remains in the membrane (Tables 2-3; Table S1). Thus, CP-stress appears to result in significantly fewer functional respiratory complexes in the plasma membrane.

**Figure 8.**
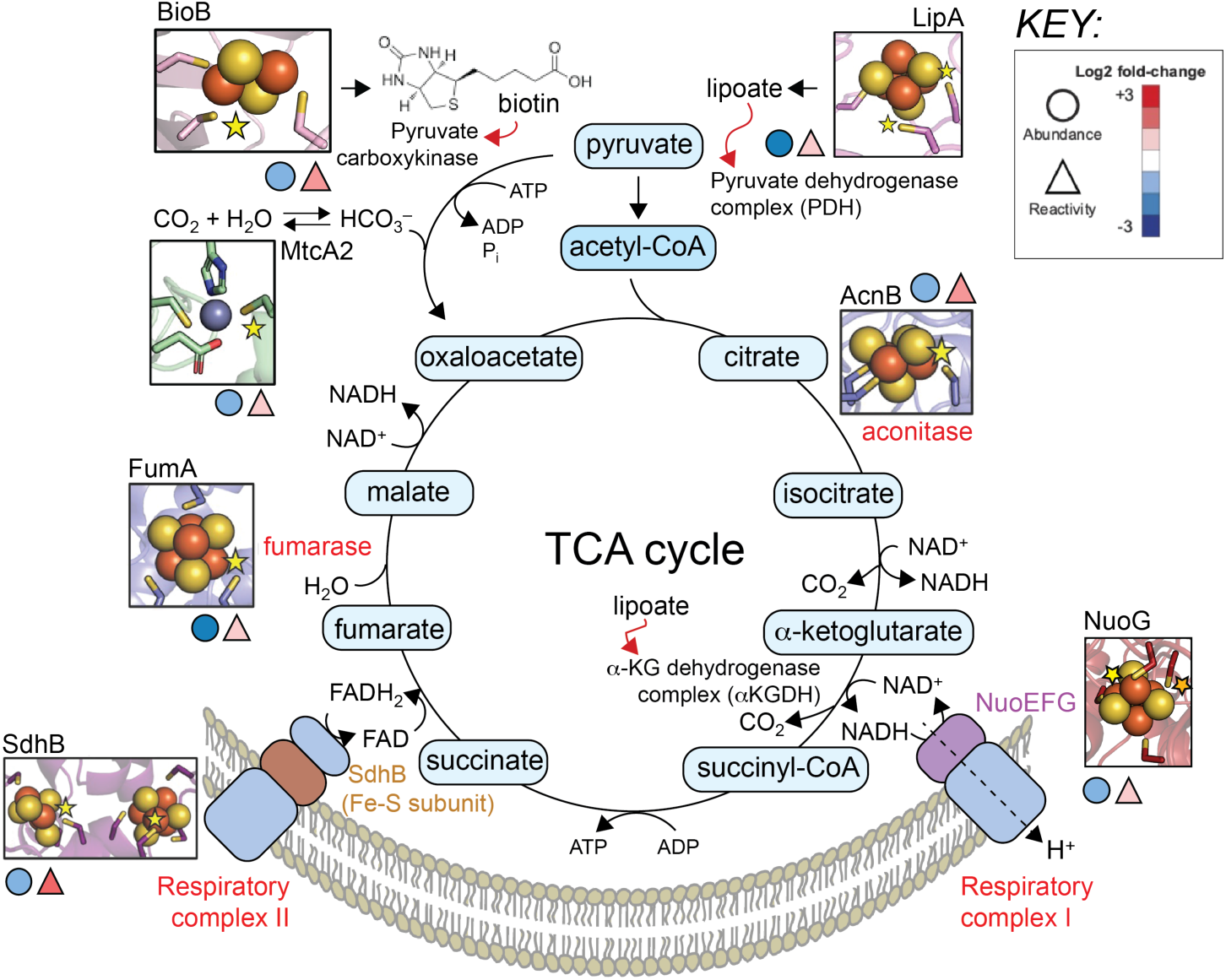
Schematic illustration of CP-induced undermetalation of enzymes involved in respiration, cofactor biosynthesis and energy transduction. The TCA cycle is coupled to the NADH dehydrogenase respiratory complex I and the succinate dehydrogenase (SDH) complex II. These electons flow to respiratory complexes III and IV to create the protonmotive force that powers ATP synthesis (not shown). Fe-S and Zn coordination sites are shown, with stars indicating the Cys that become more reactive under conditions of CP stress. Shaded circles and triangles, see key.

### Enzymes in Other Metabolic Pathways that are Negatively Impacted by CP stress

Metalloproteins that are undermetalated (Table 3; Table S1) are associated with disparate cellular processes, and a number of these Zn enzymes are associated with GTP metabolism (Figure S6). The first two enzymes of the “lower” arm of the flavin biosynthesis pathway are catalyzed by the consecutive action of two Zn enzymes, GTP cyclohydrolase II (RibA) and NADP^+^-dependent diaminohydroxyphospho-ribosylaminopyrimidine deaminase (RibD).^30^ C55 in RibA (log_2_=0.90) and C88 in RibD (log_2_=2.72) are significantly undermetalated, with RibD among the most profoundly impacted Zn enzymes in the cell (Table 3). These findings are consistent with previous work that showed that *A. baumannii* employs an alternative 3,4-dihydroxy-2-butanone 4-phosphate synthase (DHBPS; RibBX) that catalyzes the first step of the “upper” arm of flavin biosynthesis pathway in an effort to maintain cellular flavin levels.^30^ These data taken together suggest that CP-stress negatively impacts flavin biosynthesis with an important bottleneck in the sequential activities of RibA and RibD that occurs specifically as a result of undermetalation.

GTP is also the substrate for the Zn-enzyme GTP cyclohydrolase I (FolE) which makes the intermediate H_2_-NTP, a metabolic branchpoint in folate and the queuosine (Q)-tRNA biosynthesis pathways.^31^ A second metalloenzyme QueD2 (6-carboxy-5,6,7,8-tetrahydropterin (CPH_4_) synthase) catalyzes the rate determining step in Q-tRNA biosynthesis. Another Zn enzyme QueC, 7-deazo-7-deasaguanine synthase functions in the third step of the Q-tRNA biosynthesis pathway. FolE (C144), QueD2 (C18 from the inhibitory site 2^31^) and QueC (C190) are undermetalated, which pinpoints the sites of enzyme failure in cellular attempts to prioritize this pathway (Figure S6). These observations provide a molecular explanation for the previous finding that CP compromises flux through the Q-tRNA pathway.^31^

## DISCUSSION

In the work presented here, we employ a change in the reactivity of the metal-ligating cysteines in metalloproteins to pinpoint for the first time across any bacterial proteome the impact of CP stress on the metal occupancy of the Cys metalloproteome. We observe significant and widespread unndermetalation that potentially impacts many foundational cellular processes, from energy metabolism (Figures 4, 8; Figure S5) to GTP catabolism (Figure S6) to protein biosynthesis and homeostasis (Figures S7-S8). We take advantage of the fact that chronic CP- stress induces a significant Zn- and Fe-starvation regulatory response under our growth conditions, thus providing a means to evaluate and compare the integrity of the Zn and Fe-S proteomes in the same cells under the same stress conditions. We find that cellular Fe deficiency does not strongly track with reduced total cell-associated Fe under these conditions (Figure S1); this is also true in previously published work.^30^ This suggests a marked change in cellular Fe- speciation that Fur senses as Fe starvation; indeed, reduced occupancy and decreased levels of cell-abundant Fe-S enzymes (under metal-replete conditions) is direct evidence of a pronounced perturbation of Fe-speciation in CP-stressed cells, which is effectively Fe-sparing (Figure 7). Remarkably, the cellular fate of that spared Fe is currently unknown, but it is clearly not broadly bioavailable. Several possibilities exist, including the presence of a bacterioferritin that has lost the ability to mobilize Fe(II) under these conditions,^81^ or an an yet underdiscoved Fe-storing encapsulin^82^distinct from the sulfur-storing encapsulin that is not detectable in CP-stressed *A. baumannii* proteomes.^83^

The molecular function of ZigA during adaptation to infected host-mediated nutritional immunity^27,30^ is difficult to assess from the experiments shown here. The quantitative protein abundance data show no significant differences in the global proteome upon ZigA deletion; similarly, chemoproteomics reveal minimal changes in the net cysteine reactivity across the peptides detected, with the possible exception of several ribosomal proteins, none of which are metalloproteins. Given the anticipated functional similarities between ZigA and two previously characterized Zn metallochaperones vertebrate Zng1 and *A. baumannii* MigC,^3,72,84–85^ coupled to the fact that Zn binding to the conserved Cys-x-Cys-Cys motif strongly stimuates GTPase activity,^27,57^ we would have expected to observe an increase in the reactivity of a Zn-coordinating cysteine(s) within a ZigA client protein(s). Therefore, the absence of such a finding is surprising. One possibility is that ZigA delivers metal to a non-Cys coordination site; alternatively, ZigA may simply interact with an as yet unidentified protein partner(s) in a guanine nucleotide- and/or Zn-regulated fashion, to an interface that lacks cysteines. Clearly, other approaches will have to be taken to identify a ZigA regulatory partner. One such approach would be to globally profile histidine reactivity in CP-stressed WT vs. Δ*zigA* cells, which may well provide significantly deeper coverage specifically of the zinc metalloproteome and thus the functional role of ZigA.^86^

In general, undermetalation of the Zn proteome appears more nuanced than that of the Fe-S cluster proteome, and encompasses enzymes and proteins that vary signficantly in their relative cell abundance (Figure 7). In an effort to identify a structural feature that might predispose a Cys-Zn proteome site to metal loss under conditions of CP-induced metal limitation, we further analyzed our undermetalated Zn proteome sites and compared these sites to those that did not meet our statistical criteria for significant loss of metal as well as to Cys proteome. Zn-sites rich is cysteine sulfur ligation might be anticipated to form slightly higher affinity complexes compared to those consisting of lighter N/O atoms;^87^ in this case, tetrathiolate (S_4_) sites might exhibit higher resistance to metal loss. We find that fraction of undermetalated S_4_ Zn-sites matches their estimated percentage in the Cys-Zn proteome (≈50%). Alternatively, the proximity of Zn ligands in the primary structure may also impact Zn loss if cotranslational metal insertion in the ribosome exit tunnel contributes significantly to metalation of the Zn-Cys metalloproteome. If we consider all four Zn ligands as being “close” in the primary structure as within am ≈40 residues in the primary structure,^88^ 60% of the undermetalated sites we observe here meet this definition; however, this is close to the approximate number of sites in the Zn-Cys proteome that that are close in the primary structure. Therefore, we cannot easily identify a specific structural feature in these coordination sites that might distinguish them from Zn-sites that are unaffected by CP-stress. It will be necessary to broadly investigate the thermodynamics of metal binding, as well as the kinetics of metal dissociation to investigate this further.

We also find that the steady-state levels of these undermetalated Zn enzymes are not severely impacted, with the exception of ribosomal protein L31. L31 is expected to have very little structure once dissociated from the ribosome, which will be exacerbated by the loss of the bound Zn, making L31 an excellent substrate for cellular proteases (Figure 3E). Only three other Zn enzymes are characterized by significantly decreased abundance coupled to reduced metal occupancy, including the β-class carbonic anhydrase and two of the five undermetalated tRNA synthetases, isoleucine- and cysteine-tRNA synthetases. This general response of the Zn proteome to metal starvation contrasts with those seven undermetalated enzymes of the Fe-S proteome, all of which are reduced in steady-state abundance, relative to those that retain their Fe-S clusters. This suggests that the metalation status of the Fe-S proteome is more strongly connected to protein homeostasis (biosynthesis, degradation and post-translational modification) than is the Zn proteome, which appears largely uncoupled from proteostasis mechanisms. The reasons for this are unknown, but one possibility is that reduced Zn occupancy fails to engage an unfolded protein response, which might in turn lead to enhanced degradation and cellular turnover of these apo-enzymes relative to the Fe-S proteome. This is turn is broadly consistent with the set-point model, in that the Cys-Zn metalloproteome exists as a superposition of non-metalated and metalated pairs of enzymes, the metalation status of which appears broadly responsive to the bioavailable Zn pool.^37^ In this case, a careful analysis of the zinc binding affinities of all 29 Cys-Zn enzymes might define the Zn set-point for chronically CP-stressed cells.

The work presented here also suggests that CP-stress has many hallmarks of nutritional stress while at the same time potentially disarming the ability of cells to fully engage the stringent response as a consequence of undermetalation of key players in this process. One metabolic fate of GTP is its hyperphosphorylation to the alarmone (p)ppGpp by GTP pyrophosphatase (RelA) (Figure S6) as part of the stringent response to general nutrient starvation. RelA associates with ribosomes in an unacylated A-site tRNA conformation and contains a C-terminal regulatory Zn-finger domain (ZFD) that has been shown to stabilize the interaction with the ribosome; mutagenesis of a key Cys in the ZBD (equivalent to our hypermodified C662 in *A. baumannii* RelA) is expected to impact association with the ribosome, which would lead to a reduced stability in cells.^89^ ^89–90^ This undermetalation of RelA in CP- stressed cells is read out by enhanced reactivity of C662 thus potentially undermining cellular efforts to mount the stringent response (Figure S6).

Indeed, as found earlier,^30^ CP-stressed cells have a lower complement of ribosomes (≈60% of untreated cells) (Figure S7A); these cells are also strongly enriched in two ribosome hibernation factors, RaiA and Hpf (hibernation promoting factor), each of which binds to the 70S ribosome and inactivates translation via distinct mechanisms (Table 2; Figure S7B).^91–92^ 70S ribosomes purified from CP-stressed cultures appear enriched for Hpf; in addition, these 70S particles are also enriched for small protein B (SmpB), despite no change in its cellular abundance (Figure S7A; Document S2). SmpB, in complex with alanyl-charged tm (transfer-messenger) RNA mediates *trans*-translation, the major rescue process for stalled ribosomes that result from a defective mRNA (“non-stop”), or an empty (deacylated) A-site (“no-go”) that results from amino acid limitation (Figure S7B), in a process that is functionally parallel to (p)ppGpp synthesis.^93–94^ Interesting Ala-tRNA synthetase is one of five highly cell abundant tRNA synthetases that is undermetalated in CP-stressed cells, which may in turn compromise this rescue process (Figure S7C). Thus, 70S ribosomes from CP-stressed *A. baumannii* cells are not only lower in number, but those that are present may be in various states of maturation^95–96^ or functionality. These findings mirror those in mycobacteria where Zn depletion induces ribosome hibernation.^55^

The stringent response is employed by bacteria to combat nutrient deprivation. Mechanisms that are used to globally remodel the proteome, including the selected use of tRNA modifications (Figures S6-S7), proteostasis, sRNA regulation and RNA processing (Figure S8), may well be compromised by significant undermetalation of key enzymes that maintain these processes. These enzymes include DnaJ, a housekeeping hsp40-class molecular co-chaperone that helps to maintain the integrity of the proteome (Figure S8A), and which is among the top-most undermetalated proteins in cells (Table 3); further, the Cys in DnaJ that reports on metalation status (C167) is a ligand to a zinc site II that is essiential for the interaction with DnaK.^97^ Others include a Cys residue in the SecY-interacting motif of SecA, an ATPase motor protein that powers movement of unfolded proteins destined for the periplasm or the extracellular space through the SecYEG channel,^98^ and the Cys residues of the “Zn-link” motif of RNase E,^99^ an essential component of the RNA degradasome^100^ intimately involved in sRNA-mediated regulation of gene expression (Figure S8B).

If undermetalation of the above-indicted Zn enzymes (Figures S6-S8) even partially impacts their cellular activities, this might be expected to induce a broad remodeling of the proteome. These perturbations might in turn be expected to induce a “knock-on” effect akin to Q-tRNA and ms^2^i^6^A-tRNA modification (which requires the 4Fe-4S cluster-containing methylthiotransferase MiaB^101^ that is undermetalated in these cells; see Table 3) in transfer-RNA pools, and in the integrity of tRNA charging itself (Figure S7) under conditions of nutritional stress.^102^ In fact, some fraction of the proteome that increases in cellular abundance in the “other” category (see Figure 2) may be attributed to indirect effects like these. Additional experiments, including a comprehensive analysis of CP-stressed metalloproteome and the metabolome will provide deeper insights into metabolic re-wiring in CP-stressed cells. This work is underway.

## MATERIALS and METHODS

### Chemoproteomics and Mass Spectrometry

#### Generation of A. baumannii lysates

Calprotectin was expressed in *E. coli* and purified as previously reported.^12,16^ Wild-type (WT) and Δ*zigA A. baumannii* cells were cultured overnight as biological duplicate in a standard Tris-LB medium used previously.^30^ The overnight cultures were diluted 1:50 into 2 mL Tris-LB media and given 1.5 h to recover at 37°C. Final cultures were prepared in 50 mL total volume of fresh media in a sterile 250 mL Erlenmeyer flask and grown with vigorous aeration until an OD_600_ of 0.370 was reached in the absence or presence of 300 µg/mL CP. The cells were harvested by centrifugation at 4000 x g at 4°C for 15 min, washed in 1 mL of 1 x PBS pH 7.4, and then immediately frozen at –80°C until further analysis. Cell pellets were thawed and resuspended in 500 µL lysis buffer (PBS, pH 7.4 + 0.1% NP-40), lysed by three rounds of sonication (10 x 1 s pulses, 85% amplitude) and centrifuged at 10,000 x g at 4°C for 15 min to generate clarified lysates. Protein concentration was determined by Bradford assay (BioRad) and lysates were diluted to a concentration of 1 mg/mL with lysis buffer.

#### SP3 purification and trypsin digestion

Samples were prepared in biological duplicate for each study condition, for a total of six (6) study channels (2x untreated WT, 2x CP (300 µg/mL)-stressed WT, and 2x CP-stressed Δ*zigA* cells). For each channel, 50 µL of lysate (50 µg, 1 mg/mL) was portioned into a low-bind 1.5 mL Eppendorf tube with 50 µL of lysis buffer. Each channel was treated with of 5 µL of 10 mM iodoacetamide-desthiobiotin (DBIA) for 1 h at room temperature in the dark with intermittent vortexing every 15 min. After DBIA labeling, 15 µL of magnetic SP3 beads (1:1 hydrophobic:hydrophilic, 50 mg/mL) and 150 µL of 20 mM DTT in 98% ethanol were added to each channel followed by 10-min incubation with rotation. The beads were pelleted using a magnetic Eppendorf rack and the supernatant removed. The beads were washed once with 500 µL of 80% ethanol, followed by the addition of 100 µL of 20 mM iodoacetamide in lysis buffer and 30-min incubation with rotation in the dark. Next, 200 µL of 20 mM DTT in 98% ethanol was added with another 10-min incubation with rotation in the dark. Following pelleting and removal of the supernatant, the beads were washed twice with 500 µL of 80% ethanol. The washed beads for each channel were then incubated overnight with rotation at room temperature in 150 µL of 200 mM EPPS buffer, 1 mM calcium chloride, and 1.5 µg sequencing grade modified trypsin.

#### TMT-labeling

After trypsin digestion, 69 µL of acetonitrile and 6 µL of the corresponding TMT-tag (20 µg/µL in acetonitrile) was added to each of the 6 channels and the reaction was allowed to incubate at room temperature with rotation for 60 min. The reaction was then quenched by the addition of 35 µL of 5% hydroxylamine in water, followed by incubation at room temperature with rotation for 10 min. The 6 channels were combined into one low-bind 2 mL Eppendorf tube and dried on a speed-vac. The combined peptide/bead sample was then resuspended in 500 µL of 10% formic acid in water and the supernatant removed from the beads on a magnetic rack. The supernatant was then desalted with a 360 mg Sep-Pak per manufacturer’s directions. The peptide sample was eluted in a final volume of ≈1 mL into a low-bind 2 mL Eppendorf tube, with 250 µL of this elution transferred to a second low-bind 2 mL Eppendorf tube for subsequent off-line fractionation (see below “Off-line fractionation…”) and protein abundance analysis. The remaining ≈750 µL in the first Eppendorf tube was subjected to enrichment on streptavidin beads (see below “Streptavidin enrichment…”) and analyzed for cysteine reactivity. Both of the samples were dried to completeness overnight on a speed-vac.

#### Streptavidin enrichment for cysteine reactivity analysis

The combined sample (1st Eppendorf tube from the “TMT labeling” section) was resuspended in 1 mL of 100 mM HEPES, pH 7.4. Addition of 50 µL of pre-washed high-capacity streptavidin agarose beads as a 50% slurry to the samples was followed by incubation for 3 h with rotation at room temperature. The bead mixture was transferred to a Ultrafree-MC centrifugal filter (hydrophilic PTFE, 0.22 µm pore size) and centrifuged at 800xg for 30 s to remove the supernatant. The beads were subsequently washed with 3 x 300 µL of 100 mM HEPES, pH 7.4, with 0.05% NP-40, 3 x 300 µL of 100 mM HEPES, pH 7.4, and 3 x 300 µL of water. DBIA-labeled peptides were eluted from the beads with 3 x 300 µL additions of 80% acetonitrile, 0.1% formic acid (1 x 20 min incubation at room temperature, 1 x 10 min at room temperature, and 1 x 10 min at 72° C). The combined elution was dried on a speed-vac, before being resuspended in 25 µL low pH buffer A (95% H_2_O, 5% acetonitrile, 0.1% formic acid). Peptide concentration was determined by quantitative peptide assay kit (Pierce) as per the manufacturer’s instructions and the sample diluted to 100 ng/µL with low pH buffer A.

#### Off-line fractionation for protein abundance analysis

The combined sample (2nd Eppendorf tube from the “TMT labeling” section) was resuspended in 500 µL high pH buffer A (95% H_2_O, 5% acetonitrile, 10 mM ammonium bicarbonate) and loaded onto a manual injection loop connected to an Agilent 1100 Series HPLC. Peptides were separated on a 25 cm Agilent Extend-C18 column using a 60 min gradient from 20-35% high pH buffer B (10% H_2_O, 90% acetonitrile, 10 mM ammonium bicarbonate). Fractions were collected using a Gilson FC203B fraction collector into a 96 deep-well plate (0.6 min/well). Subsequent concatenation of every sixth well resulted in six pooled fractions that were dried by speed-vac and then resuspended in 50 µL of low pH buffer A. Peptide concentration was determined by quantitative peptide assay kit (Pierce) per manufacturer’s instructions and each fraction was diluted to 100 ng/µL with low pH buffer A.

#### LC-MS/MS of peptide samples

Mass spectrometry-based proteomic analysis of each sample/fraction was performed as technical replicates, triplicate for cysteine reactivity and duplicate for protein abundance. LC-MS/MS was performed on an Orbitrap Exploris 240 mass spectrometer running Xcalibur v4.4 (Thermo Scientific) coupled to a Dionex Ultimate 3000 RSLCnano system. Each sample (5 µL, 500 ng peptide) was injected directly onto an Acclaim PepMap 100 loading column. Peptides were eluted onto an Acclaim PepMap RSLC and separated with a 2 h gradient from 5% to 25% of Buffer B (20% H_2_O, 80 % MeCN, 0.1% formic acid) in Buffer A (100% H_2_O, 0.1% formic acid) at a flow rate of 0.3 µL/min. The spray voltage was set to 2.1 kV. One full MS1 scan (120,000 resolution, 350-1800 *m/z*, RF lens 65%, AGC target 300%, automatic maximum injection time, profile mode) was obtained every 2 s with dynamic exclusion (repeat count 1, duration 20 s), isotopic exclusion (assigned), and apex detection (30% desired apex window) enabled. A variable number of MS2 scans (30,000 resolution) were obtained between each MS1 scan based on the highest precursor masses, filtered for monoisotopic peak determination, intensity (5E4), and charge state (2–6). MS2 analysis consisted of the isolation of precursor ions (isolation window 0.7 *m/z*) followed by higher-energy collision dissociation (HCD, collision energy 36%).

#### Database search and quantification

The tandem MS data was analyzed by the Thermo Proteome Discoverer V2.4 software package and searched using the SequestHT and Percolator algorithms against a UniprotKB database (www.uniprot.org) of the *A. baumannii* proteome (UP000072389 – 02/15/2024). Trypsin was specified as the protease with a maximum of two missed cleavages. Peptide precursor mass tolerance was set to 10 ppm with a fragment mass tolerance of 0.02 Da. The false discovery rate (FDR) for peptide identification was set to 1%.

Oxidation of methionine (+15.995) as well as acetylation (+42.011) and/or methionine-loss (+131.040) of the protein N-terminus were set as dynamic modifications and TMT-tag labeling (+229.163) of lysine residues and peptide N-termini were set as static modifications. For protein abundance analysis, cysteine alkylation (+57.021) was set as a static modification, while for cysteine reactivity analysis, cysteine alkylation (+57.021) and DBIA labeling (+296.185) were both set as dynamic modifications. For TMT-based quantification, a centroid tolerance of 20 ppm was applied for integration, with a co-isolation threshold of 50% and an average reporter signal-to-noise threshold of 10. Normalization of each channel was applied based on the total peptide signal of all peptides. Peptide ratio quantification was based on S/N values, while protein ratios were calculated from grouped abundances excluding cysteine containing peptides. P-values were calculated by ANOVA of individual peptide or protein abundances. Annotation of cysteine function was generated from the Uniprot Protein Knowledgebase (UniProtKB).

### Fe-S Cluster and Zn Proteome Predictions

For the prediction of the Fe-S proteome, we used the MetalPredator server^58^ to scan the *A. baumannii* ATCC17978-VU proteome with default parameters. This resulted in a list of 92 putative Fe-S cluster proteins (Document S1). For the prediction of the Zn proteome, we used the approaches previously employed,^30,58^ creating two libraries of Hidden Markov Model (HMM) profiles, one based on known Zn-binding Pfam domains, and another based on specific Zn-binding structural motifs. The former library was created by combining domains with a known 3D structure and domains without a known structure but annotated as Zn-binding. The latter library was obtained by fragmenting Zn-binding sites from existing protein structures, while excluding non-physiological sites. We then used these two libraries to scan the *A. baumannii* ATCC 17978-VU proteome with the hmmscan tool,^103^ filtering out proteins matching HMM profiles that did not conserve the Zn-binding residues. This procedure yielded a list of 203 putative Zn proteins, which was integrated with data from the UniProt database resulting in a final list of 225 putative Zn proteins (Document S1).

### Multireaction Monitoring for Quantitation of ZigA

#### Generation of ^15^N-ZigA spiked A. baumannii lysates

Pipette stabs from a frozen glycerol stock of ATCC 17978-VU were transferred into triplicate cultures containing 1 mL Tris-LB and grown with shaking at 37°C for 16 h. Overnight cultures were diluted 1:50 into 1 mL fresh media and given 1 h to recover in the 37 °C incubator. Recovery cultures finally diluted 1:500 into 10 mL Tris-LB with and without 300 µg/mL calprotectin. Cells were collected once cultures reached an OD600 of 0.4, washed with 1 mL wash buffer (25 mM Tris pH 7.5, 100 mM NaCl), and stored at –80 °C until use.

Pellets were resuspended in 600 µL lysis buffer (25 mM HEPES pH 7.4, 150 mM NaCl, 2.5 mM TCEP, EDTA-free protease inhibitor) and transferred to lysing matrix B tubes, which were then lysed on an Omni Bead Ruptor 12 homogenizer by performing a standard lysis protocol (6 m/s, 3 min on, 1.5 min off, 30 s total) twice. Lysates were centrifuged at 20,000 x g for 10 min and the supernatants were decanted into clean 1.5 mL Eppendorf tubes. This step was repeated in order to remove any residual beads and insoluble material. ^15^N-*Ab*ZigA was spiked into each lysate replicate with the appropriate amount added based on preliminary lysate analysis and the standard curve that was generated from ratiometric standards. Total protein content in each sample was then measured by Bradford assay.

#### Preparation of ^15^N/^14^N-AbZigA Ratiometric Standards

Uniformly labeled ^15^N-and unlabeled-*Ab*ZigA were purified as previously described.^57^ Seven quantitation standards were generated by mixing calculated mass ratios of ^15^N:^14^N protein in triplicate ranging from 0 to 1.0. The normalized ^15^N:^14^N mass ratios were plotted against their normalized ^15^N:^14^N signal ratios for multiple tryptic peptides to generate standard curves.

#### Trypsin Digest and Peptide Desalting

20 µg of protein (standard or lysate) was transferred to a clean Eppendorf tube and dried down using a SpeedVac concentrator. Pellets were resuspended in 100 µL 8M urea and 100 mM ammonium bicarbonate solution. Then, 2 µL 0.5 M TCEP and 10 µL 200 mM iodoacetamide was added to each sample and stored for 1 h in the dark at room temperature for subsequent reduction and alkylation. Protein precipitation was initiated by adding 25 µL of trichloroacetic acid and continued for 1 h at –20 °C. Samples were pelleted in a SpeedVac by centrifugation for 20 min at 20,000 x g, and the supernatant was carefully removed. Proteins were washed with 500 µL of cold acetone, and the samples were dried by SpeedVac concentration. Trypsin digestion was initiated after dissolving the pellets in 100 µL 1 M urea and 100 mM ammonium bicarbonate containing 2 ng/µL sequencing-grade trypsin. Resuspensions were incubated for >16 h in a shaking incubator set to 37 °C. 5 µL 10% trifluoroacetic acid was then transferred to each digest to quench the reaction and precipitate the resulting peptides. A desalting step was performed using C18 Zip Tips and the peptides were eluted in 25 µL 50% acetonitrile in water and 0.1% trifluoroacetic acid.

#### LC-MS/MS Proteomic Analysis and Calculations

Samples were analyzed on an Orbitrap Fusion Lumos Tribrid mass spectrometer coupled to an Easy-nanoLC 1200. Peptides were loaded onto an Acclaim PepMap 100 C18 trap column (75 µm x 2 cm, 100 Å) in 0.1% formic acid. The peptides were separated using an Acclaim PepMapTM 100 analytical column (75 µm x 25 cm, 100 Å) using an acetonitrile-based gradient (Solvent A:0% acetonitrile, 0.1% formic acid; Solvent B: 80% acetonitrile, 0.1% formic acid) at a flow rate of 300 nL/min. A 60 min gradient was implemented as follows: 0 min, 4% B; 0-53 min, 4-26% B; 53-55 min, 26-100% B; 55-60 min, 100% B followed by re-equilibration to 4% B. The electrospray ionization was carried out with a nanoESI source at a 260 °C capillary temperature and 1.8 kV spray voltage. The mass spectrometer was operated in data-dependent acquisition mode with mass range 400 to 2000 *m/z.* The precursor ions were selected for tandem mass (MS/MS) analysis in Orbitrap with 3 s cycle time using HCD at 35% collision energy. Intensity threshold was set at 2e5. The dynamic exclusion was set with a repeat count of 1 and exclusion duration of 60 s. The abundances of unlabeled and ^15^N-labeled versions of peptides were manually determined using the Qual Browser program within Xcalibur.^15^N:^14^N signal ratios of tryptic peptides from spiked lysates were plotted against their respective standard curves to determine the mass of ^14^N-ZigA present. The final number of ZigA proteins per cell was calculated under the assumption that an OD600 of 1 = 8×10^8^ cells.

### Label-free Proteomics Analysis of *A. baumannii* Soluble Cell Lysates

Preparation of *Ab* lysates, along with downstream digestion and peptide desalting, followed the same procedures as previously described.^30^ The LC-MS analysis was conducted in a similar manner to the preceding section. The resulting data were searched against an *Acinetobacter baumannii* database (Uniprot UP000094982, 3,780 entries, with the database downloaded on 07/06/2017 from Uniprot) in Proteome Discoverer 2.5. Carbamidomethylation of cysteine residues was set as a fixed modification. Protein N-terminal acetylation, oxidation of methionine, protein N-terminal methionine loss, protein N-terminal methionine loss and acetylation, and pyroglutamine formation were set as variable modifications. A total of 3 variable modifications were allowed.

Trypsin digestion specificity with two missed cleavage was allowed. The mass tolerance for precursor and fragment ions was set to 10 ppm and 0.05 Da respectively. Peptides abundances were quantified using the Percursor Ions Quantifier node. Protein abundances were estimated from the three most abundant peptides from a given protein and those that were assigned with less than 3 unique peptides were filtered out of the ranked abundance analysis.

### Preparation and Characterization of Fe-S Cluster-loaded *Ab*AcnA and *Ab*AcnB

#### Cloning, induction, and purification

A1S_0558 (AcnA) and A1S_2126-2128 (AcnB) were amplified from *A*. *baumannii* ATCC17978-VU genomic DNA and cloned into the pHis-parallel1 expression vector at the *Nco*I recognition site using isothermal assembly. Constructs were designed to include a 6X His-tag followed by a TEV cleavage sequence. The resulting plasmids were transformed into BL21(DE3) competent cells by heat shocking for 45 s at 42 °C. Surviving cells were selected on an LB agarose plate supplemented with 100 *μ*g/mL ampicillin that was incubated at 37 °C overnight. Colonies were then picked, transferred to 1 L LB medium containing ampicillin, and grown at 37 °C while shaking vigorously. Once cultures reached an OD600 of 0.6-0.8, 1 mL of 1 mM isopropyl β-D-1-thiogalactopyranoside was added to induce heterologous expression. The temperature was then reduced to 18 °C and incubation continued overnight. Cultures were centrifuged (5,000 x g, 4 °C, 20 min), the pellets were resuspended in 40 mL lysis buffer (25 mM Tris pH 8.0, 500 mM NaCl, 2 mM TCEP, EDTA-free protease inhibitor), and then sonicated for 10 min (2 s on, 8 s off, 60% power) in an ice bath. Insoluble material and DNA were removed from lysates with addition of 10% polyethyleneimine (PEI) and centrifugation (14,000 x g, 4°C, 20 min). Recombinant AcnA or AcnB were then bound to a prepacked, pre-equilibrated 5 mL Ni-NTA HisTrap column connected to an AKTA liquid chromatography system. The column was washed thoroughly with Buffer A (25 mM Tris pH 8.0, 500 mM NaCl, 25 mM imidazole) before eluting with a 0-100% gradient to Buffer B (25 mM Tris pH 8.0, 500 mM NaCl, 1 M imidazole). Fractions containing the desired protein were pooled, concentrated, and finally injected onto a 120 mL Sephadex G75 in buffer (25 mM Tris pH 8.0, 150 mM NaCl, 1 mM EDTA, 2 mM TCEP) for additional purification. The resulting purified protein was buffer exchanged into chelex-treated metal-free buffer (25 mM HEPES pH 7.4, 150 mM NaCl, 2 mM TCEP) by dialysis to remove EDTA and stored at –80°C.

#### Loading Fe-S clusters

Protocols to reconstitute Fe-S clusters into AcnA and AcnB were adapted from previous work.^104–105^ All buffers and reagents were freshly prepared and degassed the same day. Thawed protein was transferred into an anaerobic chamber containing <1ppm O_2_, and briefly buffer exchanged into degassed metal-free buffer. Following reduction with 10 mM DTT, eight-fold molar excess of Mohr’s salt (Fe(NH₄)₂(SO₄)₂·6H₂O) was added to the protein and the reaction was allowed to stand for 5 min. The same concentration of Na_2_S was then introduced, and the solution was incubated for 1-2 h. A color change could be observed after 30 min, typically to red or brown. Excess Fe and Na_2_S was removed after passing the samples through a PD-10 desalting column in the anaerobic chamber. An observable peak at ≈420 nm by UV-Vis confirmed the presence of Fe-S clusters.

#### AcnA and AcnB activity assays

Activity assays were performed using the Abcam colorimetric aconitase kit (ab109712). Reactions were initiated by the addition of AcnA or AcnB (500 nM final conc.) into the supplied buffer containing excess isocitrate. Formation of the intermediate cis-aconitate from isocitrate was monitored as an increase in absorbance at 240 nm (ε_240nm_ = 3.6 mM^-1^ cm^-1^). For assays containing 300 µM EDTA, enzymes were preincubated with EDTA for 30 min prior to activity measurements. Calculated activities were background corrected.

### Purification of Total Membrane Fractions from *A. baumannii*

300 mL of *Ab* ± 300 µg/mL CP were grown in Tris-LB and cells collected as described above. Further purification of the membrane fraction closely followed published methods.^106^ Briefly, cell pellets were resuspended in 12.5 mL of buffer A (0.5 M sucrose, 10 mM Tris, pH 7.5), 180 µL lysozyme was added from a freshly prepared 10 mg/mL stock and stirred on ice for 2 min. 12.5 mL of 1.5 mM EDTA was then added to the mixture and allwed to mix for 7 min before centrifugation at 11,000 x g for 10 min. The supernatant was decanted and the isolated pellet was resuspended in 5 mL buffer B (0.2 M sucrose, 10 mM Tris pH 7.5) containing EDTA-free protease inhibitor. The suspension was vigorously vortexed until homogenous and aliquoted into six lysing matrix B tubes. Cells were lysed using an Omni Bead Ruptor 12 with the standard lysis protocol (30 s total, 3 s on, 2 min off, 6 m/s) performed twice at 4° C. Lysates were centrifuged to clear cellular debris and pooled into Ti70 tubes, then diluted to the top with buffer B before ultracentrifugation at 184,500 x g for 6 h. The supernatant was carefully decanted and the remaining pellet was resuspended in 2 mL isolated membrane storage buffer (10 mM Tris-HCl, pH 7.5) by dounce homogenization. Protein concentrations were measured by Bradford assay, and total membrane fractions were stored at –20 °C until further analysis.

### NADH-quinone Dehydrogenase and Succinate-quinone Dehydrogenase Assays

NADH-quinone dehydrogenase activity in purified *A. baumannii* membranes was measured by the conversion of NADH to NAD^+^, observed by a decrease in absorbance at 340 nm (ε_340nm_ = 6.22 mM^-1^ cm^-1^). Reactions were initiated with the addition of 50 µg membrane protein into storage buffer containing 200 µM NADH and were monitored for 5 min or until the substrate was fully consumed. Calculated activities were background-corrected using reactions monitored without protein. Succinate-quinone dehydrogenase was assayed according to a published protocol.^107^ Briefly, 10 µg of membrane protein was introduced into a mixture containing 400 µM phenazine methanosulfate (PMS), 100 µM succinate-2,6-dichlorophenolindophenol (DCPIP), and 5 mM sodium succinate in buffer. Reactions were monitored by a decrease in absorbance at 600 nm (ε_600nm_ = 20.7 mM^-1^ cm^-1^), indicating successful oxidation of succinate, subsequent electron transfer to the intermediate PMS, and final reduction of DCPIP. Calculated activities were background corrected using reactions monitored without protein.

#### Purification of Ribosomes from Unstressed and CP-stressed *A. baumannii*

500 mL cultures of *Ab* ±300 µg/mL CP were grown in Tris-LB as described above and cells collected, with tight-coupled ribosomes purified as described^108^ and subjected to label-free quantitation as described above.

## ASSOCIATED CONTENT

**SI – Supporting Information**

The Supporting Information is available free of charge at https://pubs.acs.org/doi/10.1021/acschembio.xxxxx

Supporting Information including Supplementary Table S1, Supplemenatry Figures S1-S8 and Supplemental References (PDF); Supplementary Documents S1, S2 and S3 (as Excel files), which contain metalloproteome predictions (Document S1), all chemoproteomics data (Document S2) and all label-free proteomics data (Document S3).

## AUTHOR INFORMATION

Complete contact information is available at: https://pubs.acs.org/doi/10.1021/acschembio.xxxxx

### Author contributions

M. K. O. carried out the analysis and biochemical validation of the chemoproteomics data acquired by D. W. B. under the supervision of E. W. on biological samples provided by M. K. O. M. K. O. performed the all label-free proteomics experiments in collaboration with J. C. T. and P. V. C. J. M. C. and E. P. S. provided the Δ*zigA* strain used in this sudy and T. A. and W. J. C. provided the calprotectin used in this study. M. K. O. and D. P. G. wrote the manuscript with editorial input from all other authors.

### Notes

The authors declare no competing financial interests with the work reported here.

## ACKNOWLEDGEMENTS

We gratefully acknowledge support by the US National Institutes of Health grants R01 AI101171 (E. P. S, W. J. C. and D. P. G.), R35 GM118157 (D. P. G.), R01 AI127793 (W. J. C), and R35 GM134964 (E. W.) and US National Science Foundation (MCA 2122902 to P. V. C.). M. K. O. was supported by a graduate fellowship provided the Training Program in Quantitative and Chemical Biology (QCB) at Indiana Univerrsity (T32 GM131994).

