## Supplementary Information for "Exploring metalloproteome remodeling in calprotectin-stressed *Acinetobacter baumannii* using chemoproteomics"

**Table of Contents**


| Item | Title | Page |
|---|---|---|
| Table S1 | Secondary list of Zn or Fe-S cluster-harboring proteins and their metal-coordinating Cys residues | 2 |
| Table S2 | Expanded coverage of additional cysteines detected at undermetalated metal cofactor sites. | 3 |
| Figure S1 | ICP-MS analysis of the growth media and cell lysates for total metal concentrations | 4 |
| Figure S2 | Quantitation of the number of ZigA molecules per *A. baumannii* cell | 5 |
| Figure S3 | Change in cellular abundance of all proteins associated with the biosynthesis and utilization of the three major siderophores in *A. baumannii* ATCC 17978-VU | 5 |
| Figure S4 | *A. baumannii* aconitase characterization | 6 |
| Figure S5 | Respiratory complex I and complex II activities in unstressed and CP-stressed cells | 7 |
| Figure S6 | Schematic illustration of the undermetalation of Zn enzymes associated with GTP metabolism under conditions of CP stress in *A. baumannii* | 8 |
| Figure S7 | Ribosome features and functionality in CP-stressed cells. | 9 |
| Figure S8 | Schematic illustration of the undermetalation of enzymes associated with the proteostasis machinery and mRNA processing upon CP stress | 9 |
| | Supplemental References | 10 |

**Table S1**. Secondary list of selected Zn or Fe-S cluster-harboring proteins and their metal-coordinating Cys residues. These proteins show increased reactivity in CP-stressed wild-type (WT+CP) vs. WT untreated control cells ($log_2$=0.64-0.99; $\chi^2 \geq 9$; corresponding to a fold-change in Cys reactivity of 1.5 to <2.0).

| ***Uniprot Entry*** | ***KEGG Entry*** | ***Description*** | ***Gene Name*** | ***Modification Position (Cys)*** | ***Net Cys Ratio (log2) WT+CP v WT*** | ***Net Cys Ratio (log2) ΔzigA+CP v WT*** | ***Metal Cofactor*** |
|---|---|---|---|---|---|---|---|
| A0A1E3M9N2 | A1S_0403 | Ribonuclease E | *rne* | 404 | **0.96** | **0.78** | Zn |
| A0A077GGS3 | A1S_1025 | Iron-sulfur protein | *rnfB* | 102 | **0.94** | **0.96** | Fe-S cluster |
| V5VHI8 | A1S_0168 | Peptide deformylase | *pdf* | 92 | **0.93** | **1.05** | Fe |
| V5V9J6 | A1S_3407 | GTP cyclohydrolase-2 | *ribA* | 55 | **0.90** | **1.01** | Zn |
| A0A5P1UI52 | A1S_3029 | tRNA-2-methylthio-N(6)-dimethylallyladenosine synthase | *miaB* | 77 | **0.86** | **0.98** | Fe-S cluster |
| A0A5P1UKT1 | A1S_0020 | Isoleucine—tRNA ligase | *ileRS* | 931 | **0.76** | **0.57** | Zn |
| V5VAD3 | A1S_2714 | Succinate dehydrogenase iron-sulfur subunit | *sdhB* | 62 | **0.70** | **0.82** | Fe-S cluster |
| A0A1E3M509 | A1S_0713 | Quinolinate synthase | *nadA* | 209 | **0.66** | **0.56** | Fe-S cluster |
| A0A098SGG9 | A1S_0757 | NADH-quinone oxidoreductase | *nuoG* | 263 | **0.65** | **0.61** | Fe-S cluster |
| V5VGK9 | A1S_0502 | 4-hydroxy-3-methylbut-2-en-1-yl diphosphate synthase (flavodoxin) | *ispG* | 304 | **0.65** | **0.67** | Fe-S cluster |
| V5VDU6 | A1S_1631 | Iron-sulfur cluster assembly scaffold protein IscU | *iscU* | 37 | **0.64** | **1.25** | Fe-S cluster |
| A0A059ZHD8 | A1S_0288 | DNA-directed RNA polymerase subunit beta | *rpoC* | 93 | **0.64** | **0.52** | Zn |

**Table S2.** Expanded coverage of additional cysteines detected at undermetalated metal cofactor sites. All cysteine residues that reside in an Fe-S cluster or Zn binding site that exhibit a net $\log_2$-fold change in cysteine reactivity ≥ 0.5 in aat least one condition.

| ***Uniprot Entry*** | ***KEGG Entry*** | ***Description*** | ***Gene Name*** | ***Modification Position (Cys)*** | ***Net Cys Ratio (log2) WT+CP v WT*** | ***Net Cys Ratio (log2) ΔzigA+CP v WT*** | ***Metal Cofactor*** |
|---|---|---|---|---|---|---|---|
| V5V8K0 | A1S_3443 | Chaperone protein DnaJ | *dnaJ* | 164 | **3.00** | **2.65** | Zn |
| V5V8K0 | A1S_3443 | Chaperone protein DnaJ | *dnaJ* | 167 | **1.44** | **1.10** | Zn |
| V5V8K0 | A1S_3443 | Chaperone protein DnaJ | *dnaJ* | 200\|203 | **0.39** | **0.50** | Zn |
| A0A086HW53 | A1S_0531 | Small ribosomal subunit biogenesis GTPase RsgA | *rsgA* | 311 | **2.37** | **2.39** | Zn |
| A0A086HW53 | A1S_0531 | Small ribosomal subunit biogenesis GTPase RsgA | *rsgA* | 319 | **2.07** | **2.03** | Zn |
| A0A086HW53 | A1S_0531 | Small ribosomal subunit biogenesis GTPase RsgA | *rsgA* | 306 | **0.72** | **0.83** | Zn |
| A0A0H4UCN7 | A1S_3619 | Zinc ribbon domain-containing protein | | 6 | **2.16** | **2.11** | Zn |
| A0A0H4UCN7 | A1S_3619 | Zinc ribbon domain-containing protein | | 9 | **1.45** | **1.51** | Zn |
| A0A1E3MBL8 | A1S_2821 | Alkylphosphonate utilization protein | | 6 | **2.10** | **2.04** | Zn |
| A0A1E3MBL8 | A1S_2821 | Alkylphosphonate utilization protein | | 9 | **1.70** | **1.56** | Zn |
| A0A077G9Y2 | A1S_0388 | Aldehyde-activating protein | | 33 | **1.39** | **1.59** | Zn |
| A0A077G9Y2 | A1S_0388 | Aldehyde-activating protein | | 77 | **1.16** | **1.39** | Zn |
| V5VFR9 | A1S_0984 | carbonic anhydrase | *mtcA2* | 50 | **1.37** | **1.19** | Zn |
| V5VFR9 | A1S_0984 | carbonic anhydrase | *mtcA2* | 106 | **1.26** | **1.21** | Zn |
| V5VGB8 | A1S_0696 | ADP-ribose pyrophosphatase | | 26\|29 | **1.29** | **1.32** | Zn |
| V5VGB8 | A1S_0696 | ADP-ribose pyrophosphatase | | 4\|7 | **1.03** | **0.97** | Zn |
| A0A1E3M9N2 | A1S_0403 | Ribonuclease E | *rne* | 401 | **1.19** | **1.12** | Zn |
| A0A1E3M9N2 | A1S_0403 | Ribonuclease E | *rne* | 404 | **0.96** | **0.78** | Zn |
| V5VAD3 | A1S_2714 | Succinate dehydrogenase iron-sulfur subunit sdhB | *sdhB* | 210\|214 | **1.79** | **1.77** | Fe-S |
| V5VAD3 | A1S_2714 | Succinate dehydrogenase iron-sulfur subunit sdhB | *sdhB* | 77 | **1.10** | **0.91** | Fe-S |
| V5VAD3 | A1S_2714 | Succinate dehydrogenase iron-sulfur subunit sdhB | *sdhB* | 62 | **0.70** | **0.82** | Fe-S |
| A0A098SGG9 | A1S_0757 | NADH-quinone oxidoreductase | *nuoG* | 235 | **1.13** | **1.26** | Fe-S |
| A0A098SGG9 | A1S_0757 | NADH-quinone oxidoreductase | *nuoG* | 263 | **0.65** | **0.61** | Fe-S |
| A0A5P1UI52 | A1S_3079 | tRNA-2-methylthio-N(6)-dimethylallyladenosine synthase | *miaB* | 40 | **1.05** | **1.29** | Fe-S |
| A0A5P1UI52 | A1S_3079 | tRNA-2-methylthio-N(6)-dimethylallyladenosine synthase | *miaB* | 77 | **0.86** | **0.98** | Fe-S |
| A0A5P1UI52 | A1S_3079 | tRNA-2-methylthio-N(6)-dimethylallyladenosine synthase | *miaB* | 111 | **0.37** | **0.61** | Fe-S |

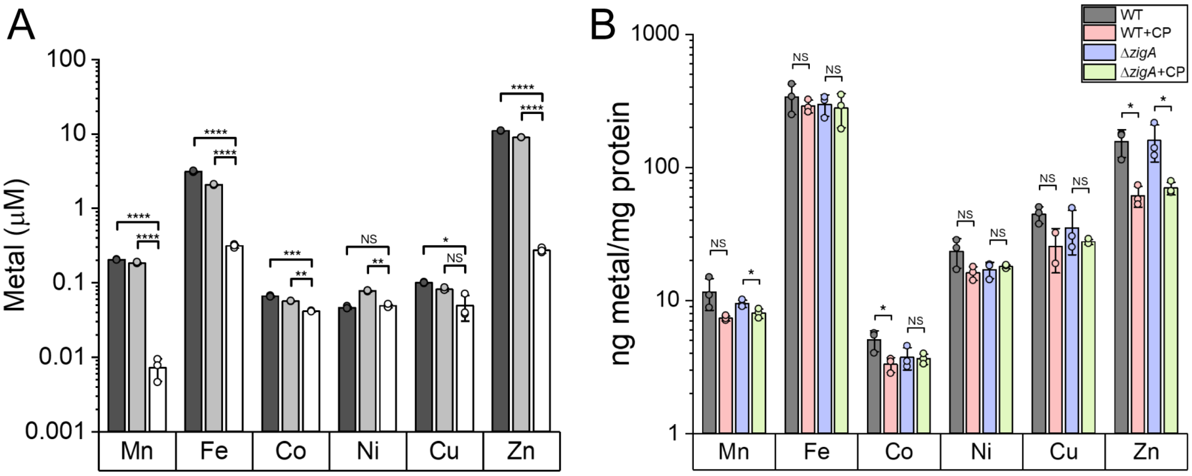


**Figure S1.** ICP-MS analysis of the growth media and cell lysates for total metal concentrations. Total metal content of LB-Tris growth media used to culture *A. baumannii* for the chemoproteomics experiments in technical triplicate (A) and of wild-type and Δ*zigA A. baumannii* cell lysates prepared in the absence and presence of 300 µg/mL CP (see *legend*) in biological triplicate (B). In panel A, total metal in the media (*black* bars) is compared with media centrifuged through a 3000 Da cut-off filter (*gray* bars) and media treated with CP and then centrifuged through the same 3000 Da cut-off filter (*white* bars) which excludes CP-bound metal. Metals are indicated from *left* to *right*. Statistical significance according to an unpaired *t*-test is indicated as follows: *, $p \leq 0.05$; **, $p \leq 0.01$; ***, $p \leq 0.001$; ****, $p \leq 0.0001$.

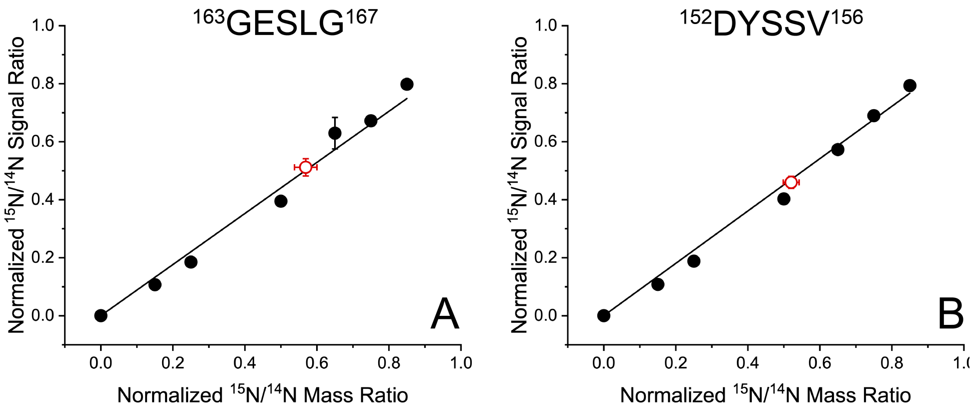


**Figure S2.** Quantitation of the number of ZigA molecules per *A. baumannii* cell using multiple reaction monitoring (MRM) with two separate peptides. (A) residues $^{163}$GESLG$^{167}$ (B) $^{152}$DYSSV$^{156}$. Calculated mass ratios suggest 410 ± 90 protein molecules per cell in CP-stressed cells, whereas ZigA is undetectable by MRM in unstressed bacterial cultures (see Methods).

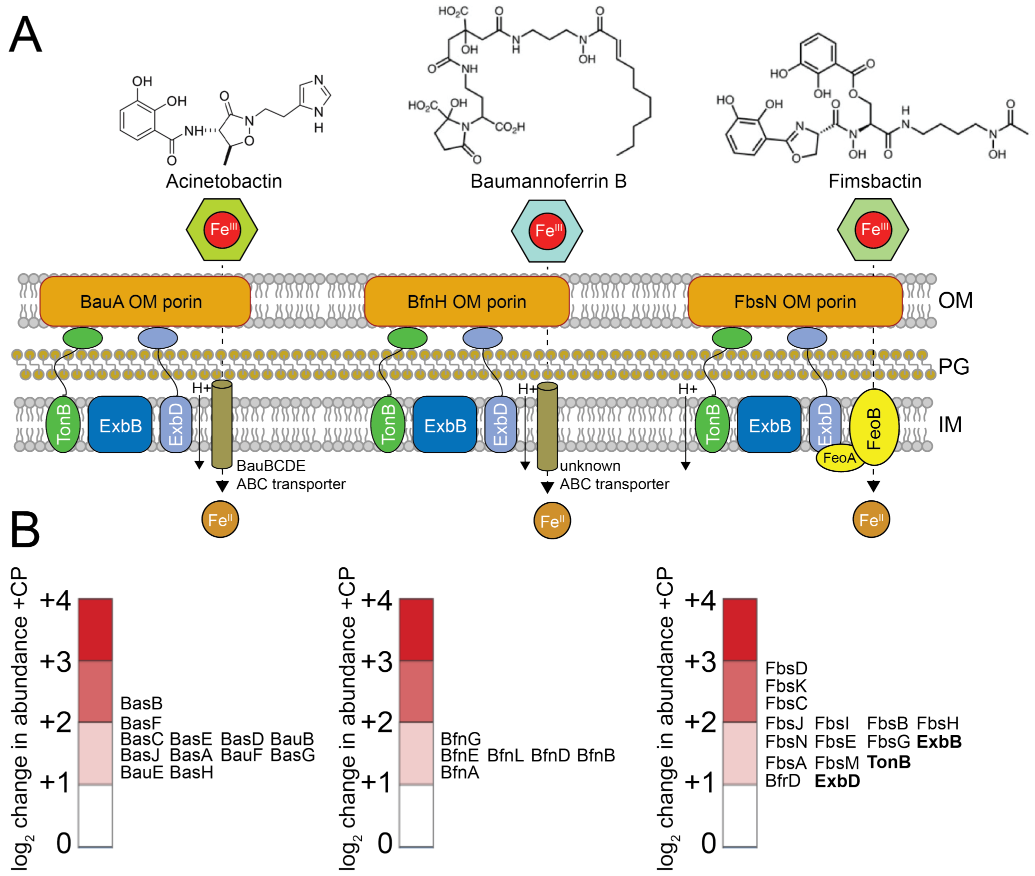


**Figure S3.** Change in cellular abundance of all proteins associated with the biosynthesis and utilization of the three major siderophores in *A. baumannii* ATCC 17978-VU. Refers to **Table 1**. (A) Acinetobactin (*left*), Baumannoferrin A and B (structure shown) (*center*) and fimsbactin (*right*). (B) Log2-fold change in cellular abundance in CP-stressed cells vs. untreated cells of the proteins and enzymes associated with the biosynthesis, efflux, and uptake of Fe(III)-siderophore complexes of acinetobactin (*left*), baumannoferrins (*center*) and fimsbactin (*right*).

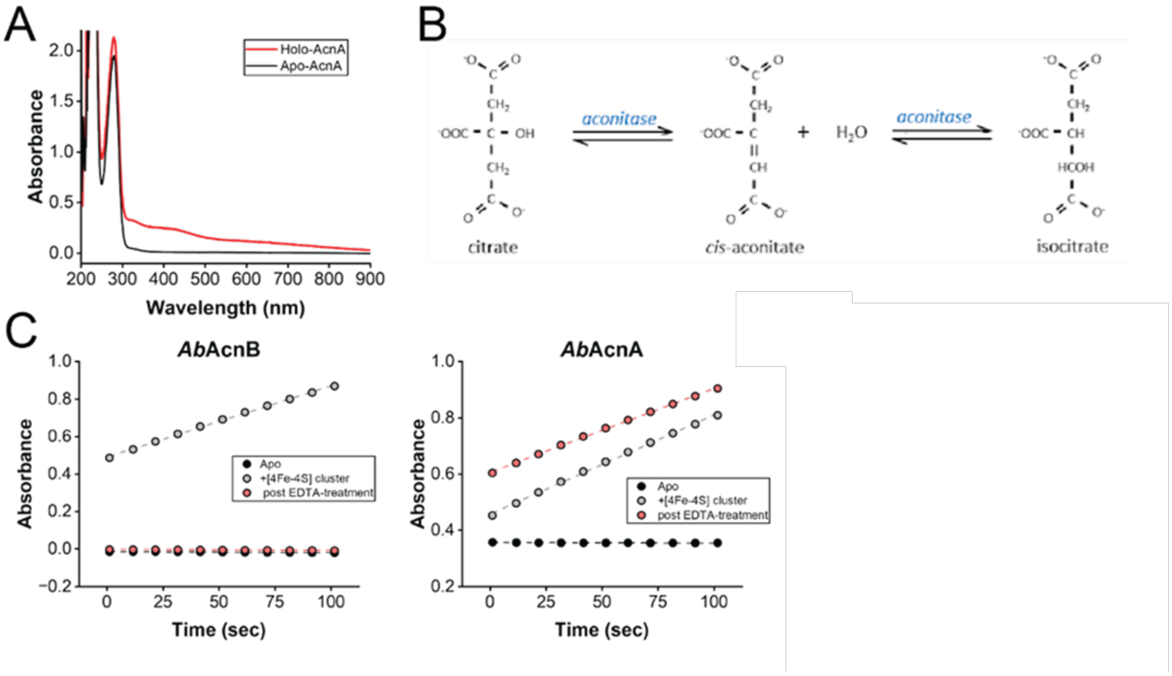


**Figure S4.** *A. baumannii* aconitase characterization. Recombinant *A. baumannii* aconitase B (AcnB) and aconitase A (AcnA) contain a 4Fe-4S cluster and very similar specific activities, but only AcnB is inactivated by EDTA treatment. (A) Absorption spectra of apo (as isolated) and 4Fe-4S cluster-reconstituted *Ab*AcnA. (B) Reaction catalyzed by aconitase, with formation of the *cis*-aconitate intermediate monitored by absorbance at 240 nm ($\varepsilon$=3600 $M^{-1}$ $cm^{-1}$). (C) Time course of the *cis*-aconitate formation for the indicated AcnB or AcnA enzymes.

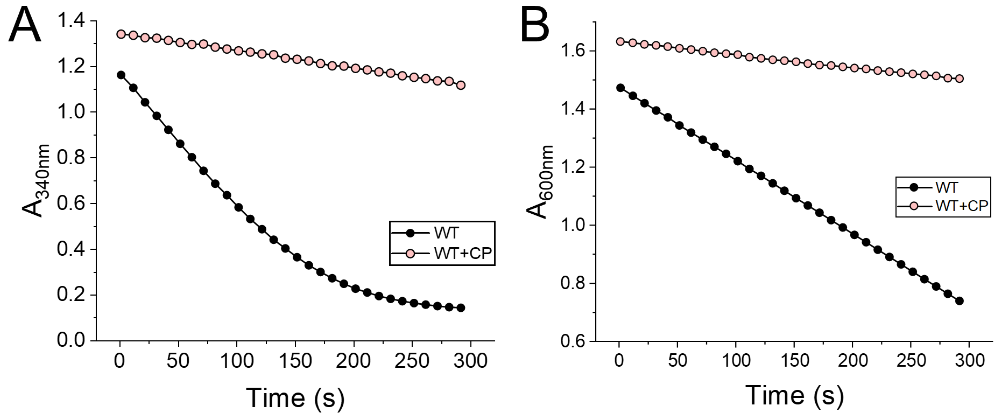


**Figure S5.** Respiratory complex I and complex II activities in unstressed and CP-stressed cells. Representative plots of (A) total membrane NADH:ubiquinone dehydrogenase (complex I) activities and (B) succinate dehydrogenase (complex II) activities in biological duplicate measured in untreated and (WT) an CP-stressed cells (WT+CP).[1]

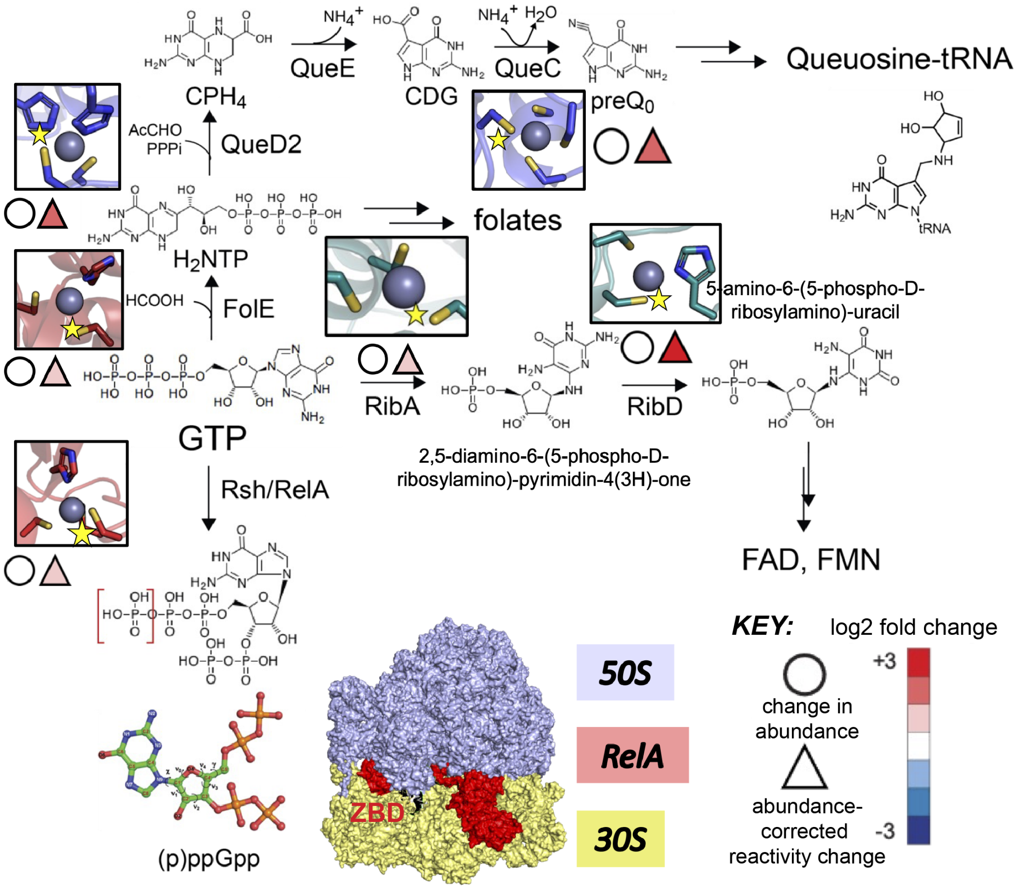


**Figure S6.** Schematic illustration of the undermetalation of Zn enzymes that function in GTP metabolism under conditions of CP stress in *A. baumannii*. Alphafold2 models of the Zn coordination sites of those enzymes that have reduced metal occupancies. The Cys residue that becomes more highly reactive in CP stress is indicated by the yellow star (see Table 3 for residue number). *Key*, color-coded change in abundance (circle) or Cys reactivity (triangle) shaded according to the log2-fold change as indicated for each metal-liganding Cys. The space-filling model of the RelA-bound *E. coli* 70S ribosome, with the $Cys_3His$ zinc finger domain (ZBD) found on the small lobe to the left bound to the A-site finger.[2]

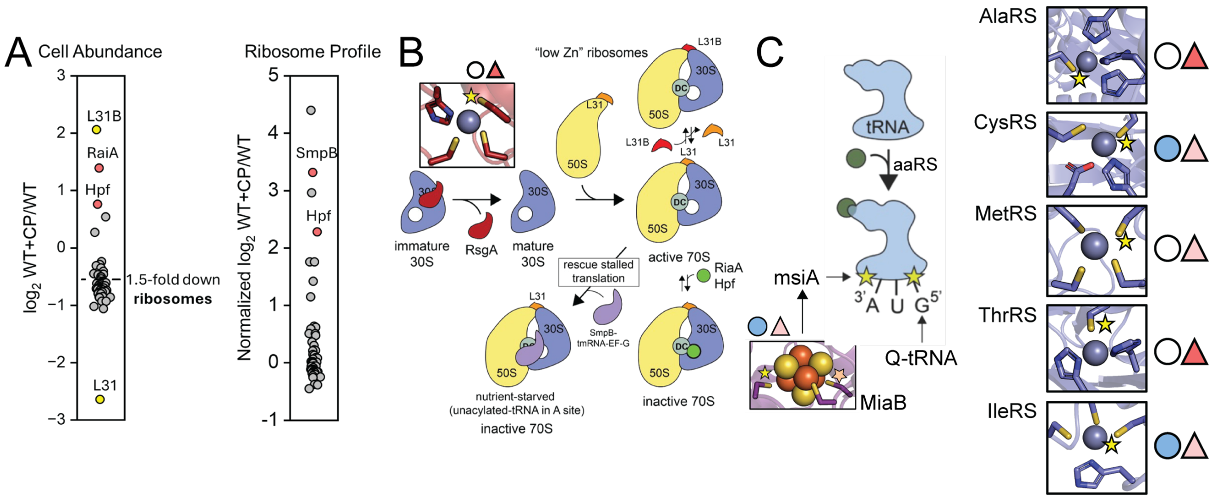

**Figure S7**. Ribosome features and functionality in CP-stressed cells. (A) *Left*, relative cell abundance of ribosomal and ribosomal associated proteins in CP-stressed vs. untreated WT cells. *Right*, normalized protein context of purified 70S ribosomes from CP-stressed vs. untreated WT cells from a label-free proteomics experiment. (B) Schematic illustration of the maturation and remodeling of ribosomes from CP-treated cells. RsgA is Zn-containing GTPase (Zn site is shown) required for maturation of the decoding center of the 30S subunit prior to assembly of the 70S particle.[3] (C) Schematic view of the tRNA charging reaction and close-up views of the Zn-sites of five tRNA synthetases (aaRS) which become undermetalated in CP-stressed cells. Also indicated are the sites of two tRNA post-transcriptional modifications, queuosine-tRNA (see Figure S6) and msiA (2-methylthio-$N^6$-dimethylallyl-A), which occurs 3' to the 3' nucleotide of selected tRNAs to enhance decoding of UNN codons.[4] The 4Fe-4S site 1 of MiaB is shown;[5] MiaB catalyzes the 2-methylthioation step. Shaded circles and triangles, see key in Figure S6 or Figure 8, main text.

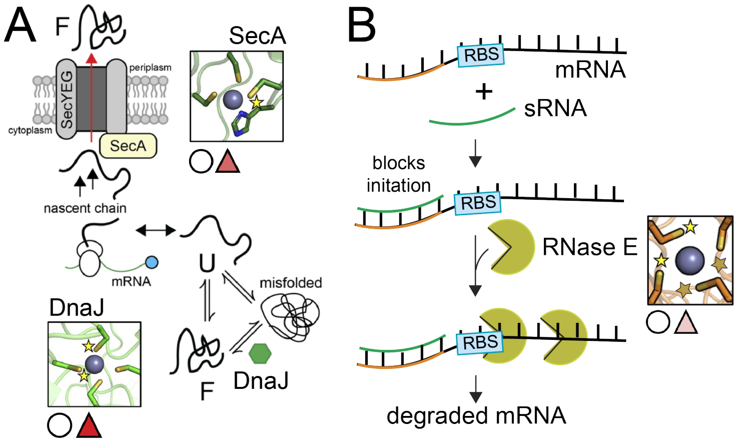

**Figure S8.** Schematic illustration of the undermetalation of enzymes associated with the proteostasis machinery and mRNA processing upon CP stress. (A) Proteostasis. F, folded protein; U, unfolded protein. (B) sRNA, small regulatory RNA, which forms a duplex with a targeted mRNAs creating a substrate for the metal-bridged dimer, RNase E. Zn coordination sites are shown, with stars indicating the Cys that become more reactive under conditions of CP stress (Cys residue numbers are provided in Table 3). Shaded circles and triangles, see key in Figure S6 or Figure 8, main text.
